# Antibiotic-induced changes to the gut microbiome attenuate neuroinflammation in a mouse model of TBI

**DOI:** 10.1101/2024.05.11.593405

**Authors:** Hannah Flinn, Austin Marshall, Morgan Holcomb, Marissa Burke, Leonardo Cruz-Pineda, Sirena Soriano, Todd J. Treangen, Sonia Villapol

## Abstract

Traumatic brain injury (TBI) induces both neuroinflammation and gut microbiome dysbiosis, yet the influence of antibiotics (ABX) on TBI-related neuropathology remains unclear. We administered a broad-spectrum oral ABX regimen to deplete the gut microbiome in single and repeated TBI mouse models. In male mice, ABX treatment significantly reduced neuroinflammation and neurodegeneration post-TBI, with no effects observed in uninjured controls. ABX also altered microbiome composition and decreased serum and fecal short-chain fatty acid levels, while intestinal damage and dysbiosis were further exacerbated by TBI severity. Notably, germ-free male mice exhibited heightened neuroinflammation and larger lesion volumes following TBI, underscoring the microbiome’s essential role in recovery. Metagenomic analyses revealed *Parasutterella excrementihominis* and *Lactobacillus johnsonii* as potential ABX-resistant taxa post-injury. These findings suggest that short-term ABX treatment may attenuate TBI-induced neuroinflammation by reshaping the gut microbiome, offering directions for microbiome-targeted therapies in TBI.

## INTRODUCTION

Traumatic brain injury (TBI) is a major contributor to disability and mortality in adults, commonly caused by military activities, vehicular accidents, or workplace incidents ^1^. TBI progresses through two distinct stages, acute and chronic ^2^, each associated with specific inflammatory responses that shape secondary injury and influence long-term recovery ^3^. While many TBI symptoms improve over time, recovery is highly dependent on factors such as the severity of prior injuries, individual differences, and the effectiveness of treatment strategies. Neuroinflammation plays a central role in driving neurodegenerative processes ^4^, and repeated TBIs can worsen this degeneration ^5^. As a result, targeting inflammation has emerged as a critical focus for therapeutic intervention.

TBI is increasingly recognized as a condition that triggers widespread changes in systemic immune activation ^6^, leading to significant intestinal inflammation ^7,8^ and disruptions in gut bacterial composition, commonly called dysbiosis ^9–14^. The gut microbiota plays a crucial role in regulating brain function through the microbiota-gut-brain axis ^7,8^, influencing key processes such as neurogenesis, cytokine release, and microglial activation ^15^. However, microbiome dysbiosis may impair these protective functions and exacerbate neuroinflammation. Investigating the complex interplay between the gut microbiome and TBI pathology could provide valuable insights into therapeutic strategies aimed at reducing neuroinflammatory damage and promoting recovery.

Short-chain fatty acids (SCFAs), such as acetate, butyrate, and propionate, are key metabolites produced by gut microbes that play important roles in immune homeostasis, energy metabolism, and anti-inflammatory responses, all supporting brain recovery^16^. A decrease in SCFA-producing and mucin-degrading bacteria and an increase in pathogenic species such as *Enterobacteriaceae* often characterize TBI-induced gut dysbiosis ^17–19^. In mouse models of TBI, injury-related stress responses have been associated with reductions in beneficial gut bacteria, including *Lactobacillus gasseri*, *Ruminococcus flavefaciens*, and *Eubacterium ventriosum* ^20^. Similarly, clinical studies have reported a loss of beneficial microbes and an increase in opportunistic pathogens such as *Clostridiales* and *Enterococcus* species following severe trauma ^21^.

Antibiotics (ABX) are routinely administered in intensive care settings to prevent infections and sepsis in patients with TBI, effectively reducing injury-related and hospital-acquired infections ^22^. Early ABX intervention has been associated with improved survival ^23^ and enhanced recovery outcomes ^24^. Preclinical studies have shown that ABX combinations ^25^, despite limited systemic absorption, can deplete gut bacteria, resulting in cognitive impairments and altered neural signaling pathways ^26^. Oral administration of broad-spectrum ABX profoundly disrupts the gut microbiome and metabolite production, leading to dysbiosis ^27^. While ABX are a cornerstone of TBI care, their impact on gut microbial composition, neuroinflammatory responses, and functional recovery remains poorly understood. Many TBI patients suffer multiple injuries over time, contributing to chronic neuroinflammation, persistent cognitive and psychiatric symptoms, and progressive neurodegeneration ^28^. Repetitive TBIs further intensify neuroinflammation and impair recovery ^29^; however, how repeated injuries shape the gut microbiome and how ABX treatment modulates neuroinflammatory outcomes in this context remains unclear.

In this study, we investigated how preexisting TBI-induced gut dysbiosis influences the response to a subsequent TBI and examined the role of ABX treatment in modulating recovery. Using a severity-based TBI mouse model, we assessed the effects of ABX in both single and repeated TBI scenarios by performing fecal microbiome amplicon and metagenomic sequencing, metabolite profiling, and neuropathological analyses in ABX-treated and control groups. ABX administration significantly altered microbial community composition and metabolite levels following repeated TBI. Notably, ABX treatment attenuated neuroinflammatory responses, suggesting that microbial depletion and shifts in antibiotic-resistant taxa may contribute to neuroprotection. These findings highlight the need to investigate further the complex interplay between ABX treatment, gut microbiota, and neurological recovery following TBI.

## RESULTS

### Successful microbiome depletion mitigates neurodegeneration after acute TBI

To investigate the effects of ABX treatment on TBI outcomes, we established a mouse model subjected to either a single (1xTBI) or double/repeated (2xTBI) CCI injury and treated with antibiotics (ABX) or vehicle (VH, water) (Figure 1A). Analysis of bacterial DNA concentration in fecal samples revealed a significant reduction in microbial load in ABX-treated groups compared to VH-treated controls at 1 day following treatment, suggesting effective microbiome depletion (Figure 1B). Histological analysis showed a significant increase in cortical lesion volume in the 2xTBI-VH group compared to the 2xTBI-ABX group. In contrast, the 1xTBI+ABX group demonstrated a protective effect, exhibiting reduced lesion volume relative to the 2xTBI+ABX group (Figure 1C-D). Motor deficits were assessed using the rotarod test, where latency to fall was measured at baseline and 1 and 5 days post-injury, showing a significant decline in the 2xTBI-VH and 2xTBI-ABX mice compared to the Sh and 1xTBI-ABX mice (Figure 1E). Quantification of TUNEL+ cells demonstrated significantly higher levels of apoptosis in the 2xTBI-VH group compared to the 2xTBI-ABX-treated group in the cortex and thalamus following injury (Figure 1F, G), suggesting that increased injury severity and chronic inflammation contribute to thalamic involvement ^30,31^. ABX treatment reduces cell death following repeated injury. Representative images confirm increased TUNEL staining in the VH-treated groups (Figure 1f1-f4, g1-g4). This finding indicates that the severity of TBI progressively affects brain regions beyond the initial site of impact over time. Thalamic neurodegeneration was observed exclusively in the 2xTBI groups, and no significant changes were detected in the 1xTBI group. These findings suggest that ABX treatment following TBI reduces gut bacterial load, lesion size, and cell death.

**Figure 1.**
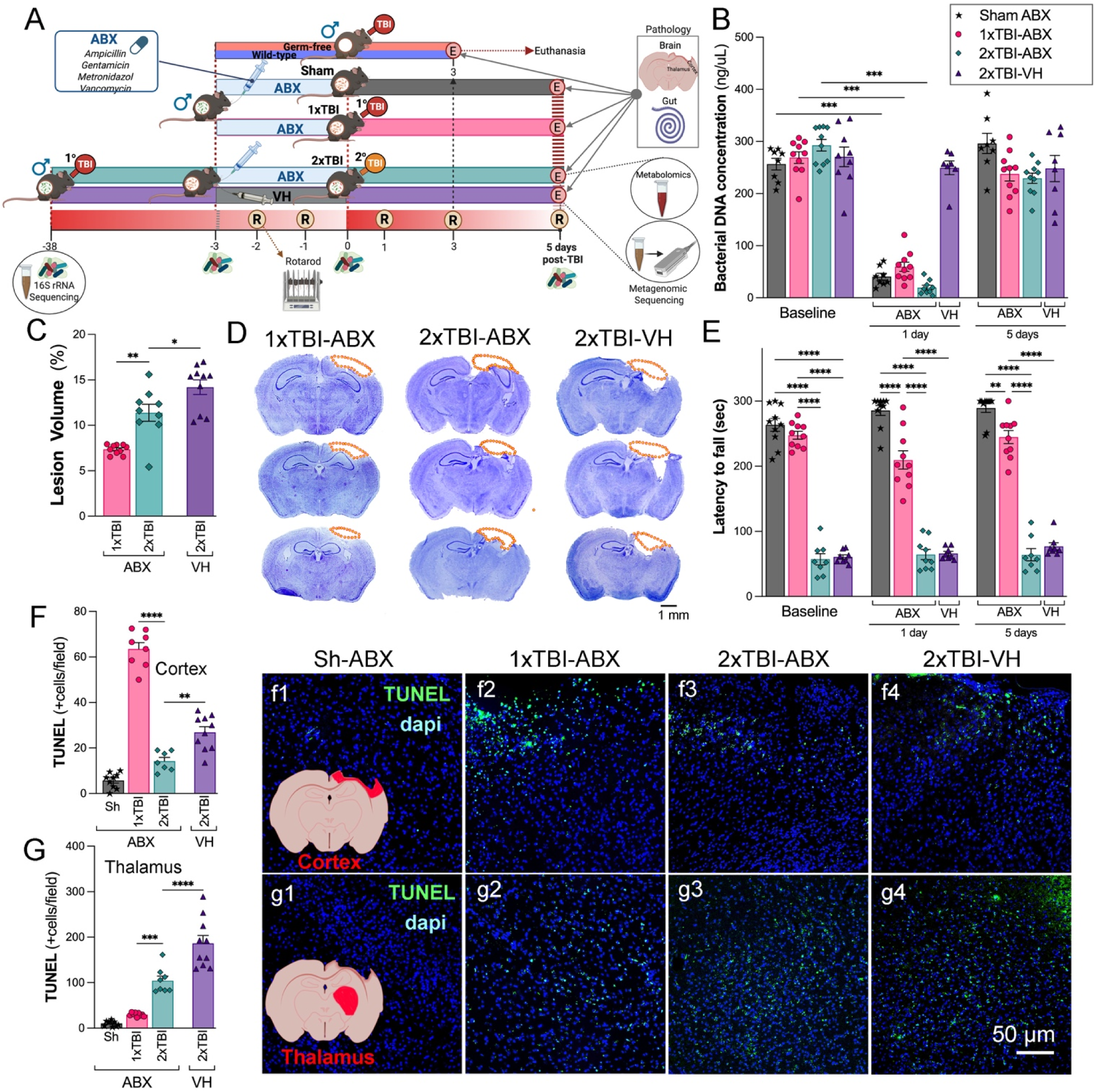
Effects of ABX-induced microbiome modulation on neuropathology in a mouse model of TBI. (A) Schematic representation of the experimental design. Mice were subjected to either sham (Sh), single TBI (1xTBI), or repeated TBI (2xTBI), with treatment groups receiving either antibiotic (ABX) or vehicle (VH). Separately, germ-free mice received a single TBI. (B) Quantification of bacterial DNA concentration across separate groups at baseline, 1 day, and 5 days post-TBI. A significant decrease in bacterial DNA concentration was observed at 1 day post-ABX, with the concentration levels returning to the baseline at 5 days post-treatment. (C) Quantification of lesion volume revealed greater damage in the 2xTBI-VH group compared to the 2xTBI-ABX group, and in the 2xTBI-ABX group compared to the 1xTBI-ABX group. (D) Representative cresyl violet-stained coronal brain sections highlighting lesion areas (outlined in orange) across experimental groups: 1xTBI-ABX, 2xTBI-ABX, and 2xTBI-VH. Scale bar = 1 mm. (E) The 1xTBI-ABX group exhibited reduced motor performance on the rotarod at both 1 and 5 days post-TBI compared to the sham-ABX group. However, their motor performance remained superior to that of both 2xTBI groups. No significant differences were observed between the 2xTBI-ABX and 2xTBI-VH groups (F, G) Quantifying TUNEL+ cells in the cortex (F) and thalamus (G) showed an increased apoptotic cell death was observed in the 2xTBI-VH groups compared to 2xTBI-ABX groups, with representative images of TUNEL (green) and DAPI (blue) staining in (f1-f4) for the cortex and (g1-g4) for the thalamus. Scale bar = 50 µm (f1-f4, g1-g4). n=7-10/group. Data is presented as a one-way ANOVA with Tukey’s post hoc test with statistical significance indicated: **p < 0.01, ***p < 0.001, ****p<0.0001.

### ABX treatment reduces microglia/macrophage activation following TBI

To assess the effects of ABX treatment on neuroinflammation following TBI, we quantified microglia/macrophage activation in the somatosensory cortex and thalamus using Iba-1 and CD68 immunostaining. The Iba-1 staining revealed a significant increase in activated microglia/macrophages in the cortex and thalamus of 2xTBI-VH mice compared to the 2xTBI-ABX groups, indicating a mitigating effect of ABX on neuroinflammation (Figure 2A, B). Similarly, CD68, a marker of activated phagocytic microglia, was significantly upregulated in the cortex and thalamus of 2xTBI-VH mice compared to ABX-treated groups (Figure 2C, D), further confirming the anti-inflammatory effects of ABX treatment (Figure 2c1–c4, d1–d4). We next evaluated P2Y12, a marker of homeostatic microglia, in the cortex and thalamus. Quantification revealed a significant reduction in P2Y12+ areas in the 2xTBI-VH group compared to the 2xTBI-ABX group, suggesting a loss of homeostatic microglial function following repeated injury and its potential to maintain microglial homeostasis (Figure 2E, F). These findings demonstrate that ABX treatment effectively reduces microglia/macrophage activation and preserves homeostatic microglial function following repeated TBI. To evaluate microglial activation and morphological changes following 2xTBI, we analyzed Iba-1 expression and conducted a Sholl analysis in the cortex and thalamus across different treatment conditions. Representative images of microglia in the 2xTBI VH group (Figure S1A-D) illustrate changes in cell morphology, including cell segmentation and skeletonization. Immunofluorescence staining for Iba-1 (Figure S1E-H) shows increased microglial density and activated morphology in the cortex and thalamus following 2xTBI, with the VH group exhibiting more pronounced activation in the cortex than the ABX-treated group. Quantitative Sholl analysis in the cortex revealed a significant reduction in microglial intersections in the ABX-treated group compared to the VH group (Figure S1I). Similarly, microglial process length was significantly decreased in the ABX group (Figure S1J), indicating a more activated state in the VH condition. However, the number of branching nodes did not significantly differ between groups (Figure S1K). In the thalamus, similar trends were observed for microglial intersections (Figure S1L), process length (Figure S1M), and branching nodes (Figure S1N), although no significant differences were detected between groups. To further assess neuroinflammatory responses, we performed *in situ* hybridization for pro-inflammatory cytokines mRNA IL-6 and TNFα in combination with Iba-1 staining (Figure S1O-R). Quantification of IL-6 mRNA+ cells showed a significant increase in the VH-treated group compared to ABX in the cortex (Figure S1S) and thalamus (Figure S1T), indicating elevated neuroinflammation in the absence of ABX treatment. Similarly, TNFα mRNA expression was significantly higher in the VH group compared to ABX in the cortex (Figure S1U), while no significant difference was observed in the thalamus (Figure S1V). These findings suggest that repeated TBI induces significant microglial activation and a robust pro-inflammatory response attenuated by ABX treatment. The results highlight the potential role of microbiome modulation in mitigating neuroinflammation and microglial reactivity following TBI. F4/80, a macrophage-specific marker, was analyzed to distinguish infiltrating macrophages from resident microglia. Quantification revealed a significant increase in F4/80+ cells in the cortex (Figure 2G) and thalamus (Figure 2H) following TBI, with the most pronounced infiltration occurring in the 2xTBI-VH group. Immunofluorescence images (Figure 2g1-4, h1-4) show more F4/80+ cell presence in TBI groups compared to the Sh, suggesting increased macrophage recruitment to the injury site. These findings demonstrate that TBI induces robust microglial and macrophage activation in both the cortex and thalamus, with repeated injury and VH treatment leading to the most pronounced neuroinflammatory response.

**Figure 2.**
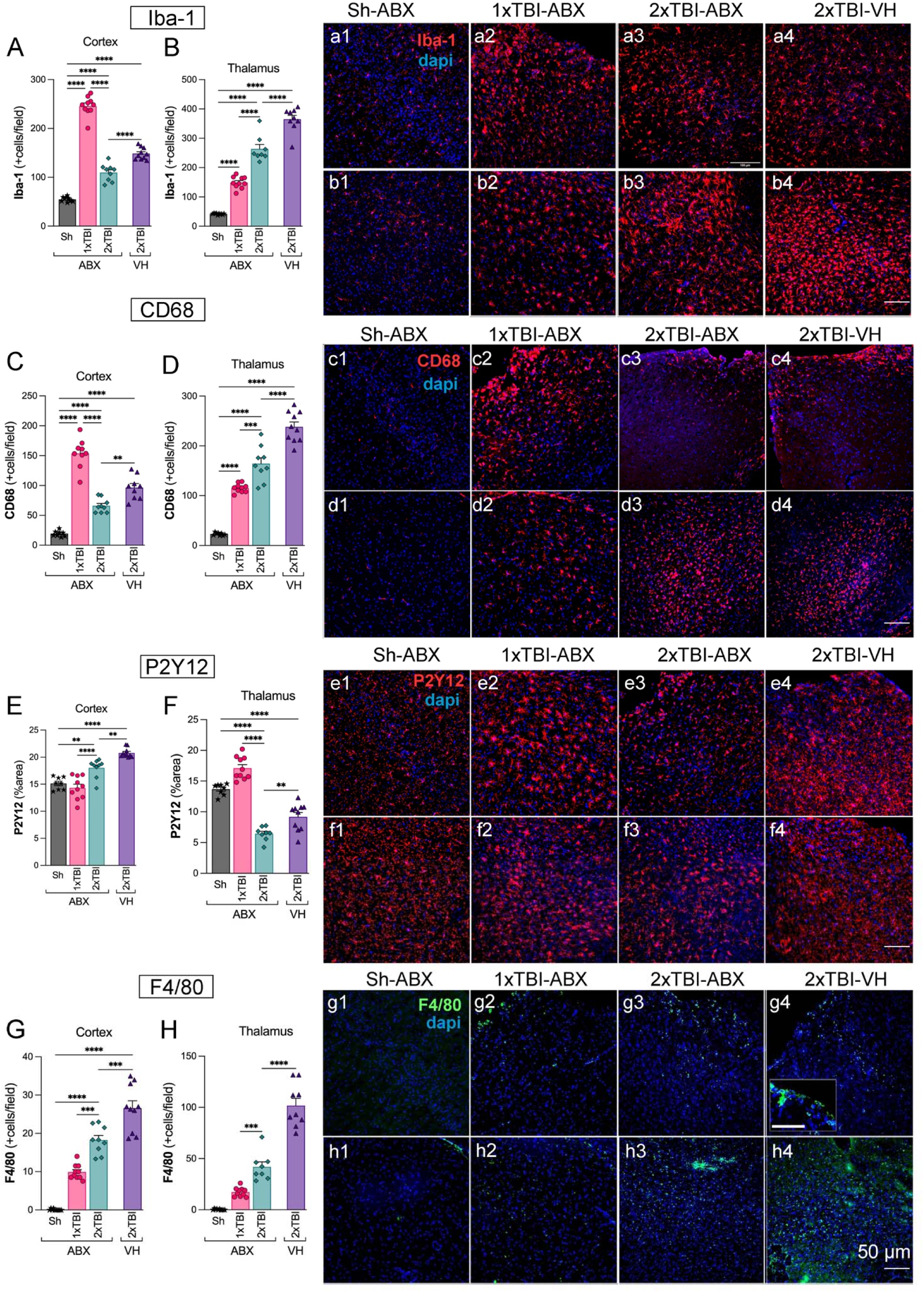
Antibiotic treatment decreases the microglia activation and macrophage infiltration in the cortex and thalamus after TBI. Quantification of Iba-1+ microglial cells in the cortex (A) and thalamus (B), showing a significant increase in microglial activation in the 2xTBI-VH group compared to ABX-treated groups. Representative Iba-1 (red) and DAPI (blue) stained images of the cortex (a1-a4) and thalamus (b1-b4) across experimental groups: Sh-ABX, 1xTBI-ABX, 2xTBI-ABX, and 2xTBI-VH. (C, D) CD68+ microglial quantification in the cortex (C) and thalamus (D) showed increased CD68 expression in the 2xTBI-VH group, indicative of a pro-inflammatory phenotype, compared to ABX-treated group and Sh-ABX. Representative CD68 (red) and DAPI (blue) images of the cortex (c1-c4) and thalamus (d1-d4) across experimental groups (E, F). Quantification of P2Y12+ microglial area in the cortex (E) and thalamus (F), showing a significant decrease in the 2xTBI-VH group, indicating loss of homeostatic microglial phenotype. ABX treatment preserved P2Y12 expression in the thalamus. Representative P2Y12 (red) and DAPI (blue) images of the cortex (e1-e4) and thalamus (f1-f4) across experimental groups. (G, H) Quantification of F4/80+ macrophages in the cortex (G) and thalamus (H), demonstrating increased infiltration in the 2xTBI-VH group compared to ABX-treated groups. Representative F4/80 (green) and DAPI (blue) images of the cortex (g1-g4) and thalamus (h1-h4) across experimental groups. Scale bar = 50 µm. n=10/group. Data is presented as a one-way ANOVA with Tukey’s post hoc test with statistical significance indicated: **p< 0.01, ***p< 0.001, ****p<0.0001.

### ABX treatment reduces astrogliosis and peripheral immune response following TBI

To assess astrocytic activation and peripheral immune cell infiltration following TBI, we analyzed the expression of GFAP and Ly6B2 in the cortex and thalamus across different experimental conditions, including Sh-ABX, 1xTBI-ABX, 2xTBI-ABX, and 2xTBI-VH. Quantification of GFAP expression, a marker of reactive astrocytes, revealed a significant increase in GFAP+ area in both the cortex (Figure S2A) and thalamus (Figure S2B) at 5 dpi. The highest levels were observed in the 2xTBI-VH group, suggesting an exacerbated astrocytic response in the absence of ABX treatment. Representative immunofluorescence images (Figure S2a1-a4, b1-b4) illustrate increased GFAP immunoreactivity in TBI groups compared to Sh, with pronounced astrogliosis in the 2xTBI-VH-treated group. To investigate peripheral immune cell infiltration, we examined the expression of Ly6B2, a marker of infiltrating neutrophils and monocytes. Quantification demonstrated a significant increase in Ly6B2+ cells in the cortex (Figure S2C) in a 2xTBI-VH group compared to 2xTBI-ABX group and between 2xTBI-ABX compared to 1xTBI-ABX in the thalamus (Figure S2D). Immunofluorescence images (Figure S2a1-a4, b1-b4) further confirm the increased presence of astrogliosis in the injured brain regions. The highest levels of Ly6B2+ cells were observed in the 2xTBI-VH group, indicating enhanced immune cell infiltration following repeated injury without ABX treatment. However, when comparing the 1xTBI-ABX and 2xTBI-ABX groups, results display increased infiltration following increased injury severity. Immunofluorescence images (Figure S2c1-c4, d1-d4) further confirm the increased presence of Ly6B2+ cells in the injured brain regions. These findings indicate that repeated TBI leads to a substantial increase in astrocytic reactivity and peripheral immune cell infiltration, with antibiotic treatment reducing the severity of these responses. The data suggest that microbiome-targeted interventions may play a role in modulating neuroinflammation following brain injury.

### Gut microbiome diversity shifts following TBI and ABX treatment

To evaluate the effects of TBI on gut microbiome diversity, we assessed alpha and beta diversity indices across different experimental groups, Sh-ABX, 1xTBI-ABX, 2xTBI-ABX, and 2xTBI-VH. Alpha diversity was evaluated using the Shannon index, which measures microbial diversity, and the Chao1 index, which estimates species richness. The Shannon index (Figure 3A) revealed a significant reduction in microbial diversity in the 2xTBI-VH groups at 1- and 5-days post-injury compared to baseline, while diversity remained relatively stable in the 1xTBI-ABX group. Similarly, the Chao1 index (Figure 3B) showed a significant decrease in species richness in the 2xTBI-ABX and 2xTBI-VH groups at 1- and 5-days post-injury compared to baseline, indicating a loss of microbial richness following injury, particularly in the VH-treated group. To further explore microbial community composition changes, principal coordinate analysis (PCoA) was performed based on Weighted Unifrac dissimilarity to quantify the variability among bacteria communities. In the Sh-ABX group (Figure 3C), microbial composition remained relatively stable across time points (baseline, 1 day, 5 days; R² = 0.677, p = 0.001), whereas in the 1xTBI-ABX group (Figure 3D), significant shifts were observed over time (R² = 0.484, p = 0.001), indicating an injury-induced alteration in gut microbiota. In the 2xTBI-ABX group (Figure 3E), the microbial composition remained significantly different from baseline at both post-injury time points (R² = 0.677, p = 0.001), suggesting a persistent effect of repeated injury. The 2xTBI-VH group (Figure 3F) exhibited the most pronounced shifts in microbial composition (R² = 0.548, p = 0.001), indicating greater gut dysbiosis without ABX treatment. These findings demonstrate that TBI induces significant changes in gut microbiome diversity and composition, with repeated injury and VH treatment leading to the most pronounced microbial dysbiosis. ABX treatment mitigates some of these changes, suggesting a potential protective effect against TBI-induced gut dysbiosis.

**Figure 3.**
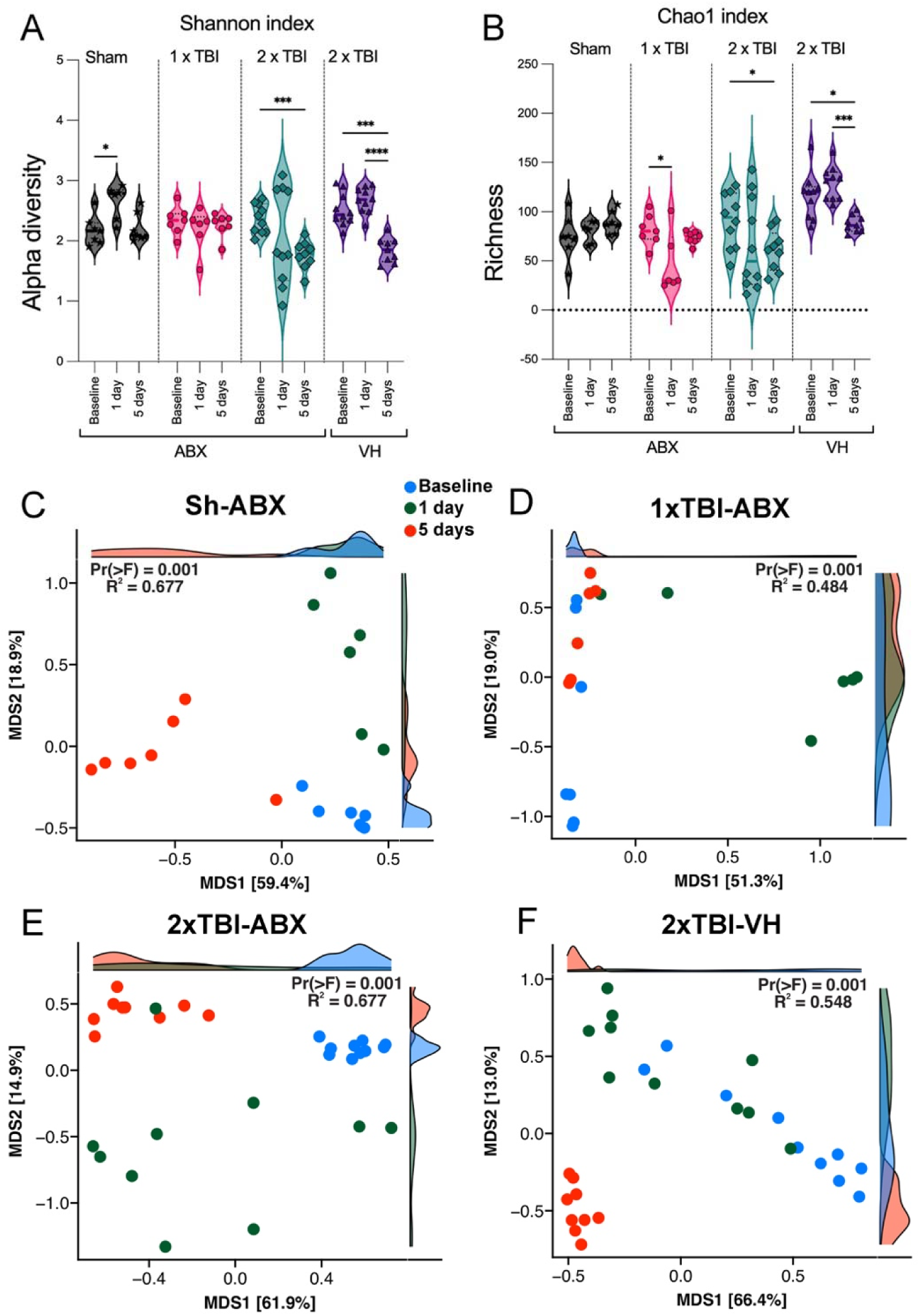
Impact of TBI and ABX treatment on gut microbiome diversity and composition. (A-B) Alpha diversity analysis of the gut microbiome in sham (Sh), 1xTBI, and 2xTBI groups treated with antibiotics (ABX) or 2xTBI vehicle (VH) at baseline, 1 day, and 5 days post-treatment (A) Shannon index, representing microbial diversity, shows a significant decrease in the 2xTBI-VH group compared to baseline, while the 2xTBI-ABX group exhibits reduced diversity over time. (B) Chao1 index, representing microbial richness, reveals a significant reduction in the 1xTBI and 2xTBI groups, with greater richness loss in the VH-treated cohort compared to the ABX-treated group. n=10/group. Statistical significance: *p < 0.05, ***p < 0.001, ****p < 0.0001. (C-F) Beta diversity analysis using principal coordinate analysis (PCoA) plots based on Weighted Unifrac dissimilarity. (C) Sh-ABX group shows significant clustering across baseline (blue), 1 day (red), and 5 days (green) post-injury (R² = 0.677, p = 0.001). (D) 1xTBI-ABX group demonstrates distinct clustering over time, indicating microbiome shifts (R² = 0.484, p = 0.001). (E) 2xTBI-ABX group exhibits significant microbial composition changes over time, similar to the Sh-ABX group (R² = 0.677, p = 0.001). (F) The 2xTBI-VH group displays the most pronounced shifts in microbial composition (R² = 0.548, p = 0.001), indicating greater dysbiosis than the ABX-treated group. These results suggest that TBI induces significant changes in gut microbiome diversity and composition, with ABX treatment mitigating some of the observed dysbiosis.

### TBI and ABX treatment induce significant changes in gut microbiome composition and metabolite production

To investigate the effects of TBI and ABX treatment on gut microbiome composition and metabolic function, we performed taxonomic profiling and SCFA analysis across experimental groups, including Sh-ABX, 1xTBI-ABX, 2xTBI-ABX and 2xTBI-VH. Longitudinal comparisons of bacterial abundance before and after antibiotic treatment revealed significant alterations in key taxa across groups. The red bars indicate increased abundance, while the blue bars signify decreases relative to baseline. We found taxa with significantly different abundance levels among the groups. In the Sh group (Figure 4A), significant increases in *Akkermansia* and *Ligilactobacillus* were observed following ABX administration, whereas decreases were seen in *Romboustia* and *Faecalibaculum*. Similarly, the 1xTBI group (Figure 4B) exhibited increases in *Akkermansia* and *Parasutterella* post-ABX treatment. In contrast, the 2xTBI-ABX group (Figure 4C) demonstrated a significant depletion of *Dubosiella* alongside an enrichment in *Parasutterella* and *Bacteroides*. 2xTBI-VH animals (Figure 4D) exhibited a distinct microbial profile, increasing *Lawsonibacte*, *Schaedlerella, Akkermansia, Bacteroides, Acutalibacter* and *Alistipes.* In addition, there is a decrease in *Clostridium*, *Lactobacillus*, and *Turicibacter*, suggesting a shift toward a dysbiotic state. A general decline was observed in the targeted SCFA quantification, with significant reductions in all SCFAs except isobutyrate in the 2xTBI-ABX group compared to the VH-treated (Figure 4E). These findings suggest that ABX treatment modulates gut microbial metabolite production, potentially influencing host metabolic and inflammatory responses by reducing SCFA-producing taxa. The overall abundance of SCFA-related GO terms hits found in long-read metagenomic data for each group normalized by read counts (Figure 4F) further supports these findings, indicating a relative shift in microbial metabolic activity between the 2xTBI-VH and 2xTBI-ABX groups. The proportion of butyrate decreased in the ABX group (36.7%) compared to the VH group (46.3%), whereas the proportions of acetate and propionate remained relatively stable between the two groups.

**Figure 4.**
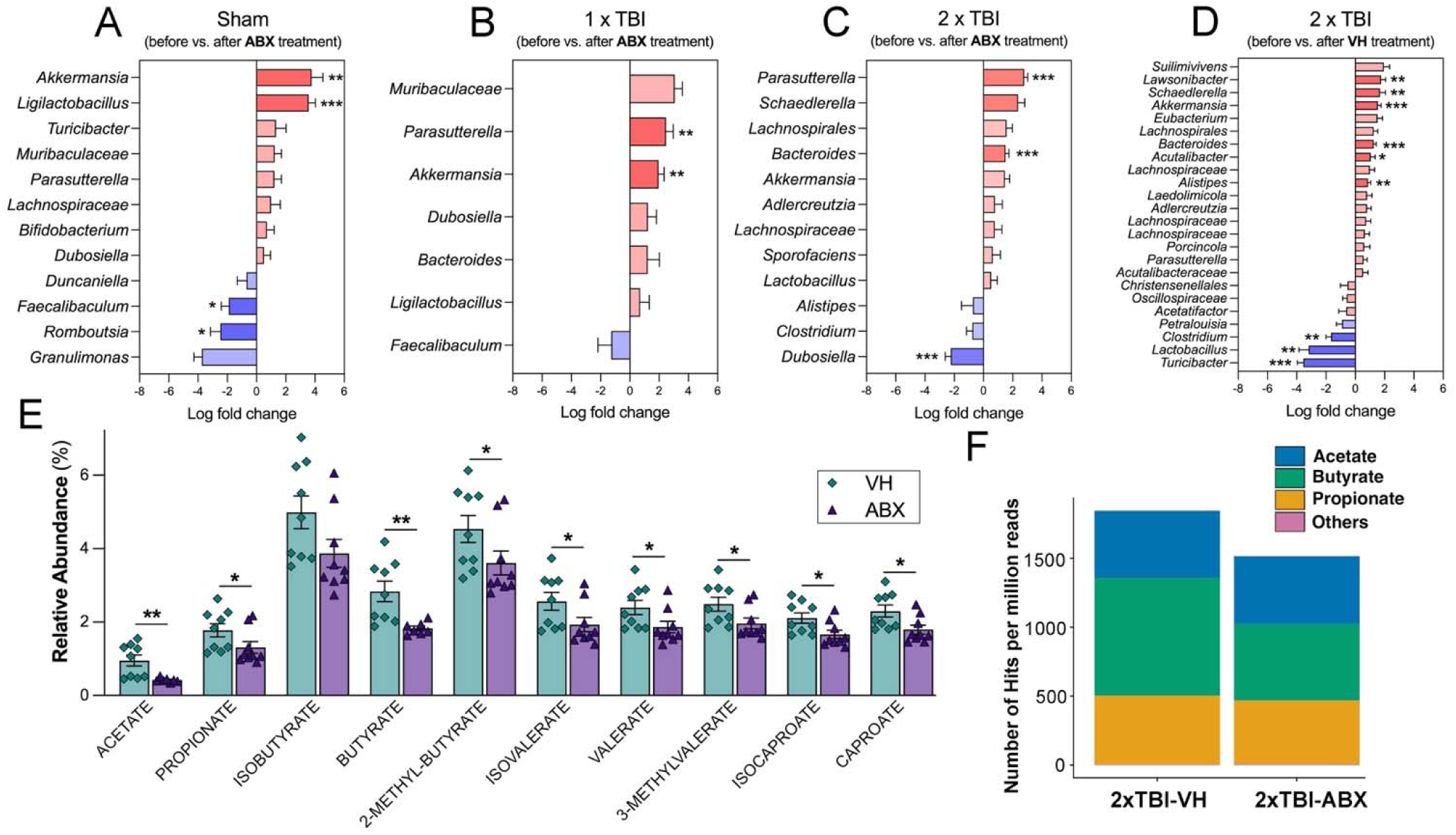
ABX treatment alters gut microbiome composition and metabolite profiles following TBI. (A-D) Log fold change in bacterial taxa before and after antibiotic (ABX) treatment in different experimental groups: (A) Sham, (B) 1xTBI, and (C) 2xTBI, with (D) showing the 2xTBI-VH group with additional microbial diversity shifts. Significant changes were observed in multiple bacterial taxa, with notable increases in beneficial taxa such as *Akkermansia* and other opportunistic bacteria post-TBI. Data are presented as log fold change with statistical significance indicated: *q < 0.05, **q < 0.01, ***q < 0.001 (ANCOMBC2). (E) Relative abundance of short-chain fatty acids (SCFAs), including acetate, propionate, butyrate, and their derivatives, in 2xTBI mice treated with vehicle (VH) or ABX. ABX treatment significantly reduced SCFA levels compared to VH. Data are presented as mean ± SEM; *p* < 0.05, p < 0.01. n=10/group. (F) Number of GO term hits per million reads corresponding to specific SCFA-producing bacterial pathways in 2xTBI-VH and 2xTBI-ABX groups. ABX treatment decreased the abundance of pathways related to acetate, butyrate, and propionate production.

### Metagenomic Insights into Gut Microbial Dysbiosis Following TBI and Antibiotic Treatment

Figure 5A highlights the gut microbiota in both 2xTBI ABX or VH-treated conditions to emphasize the unique and overlapping taxa between the groups. Notable differences were observed in the relative abundance of *Lactobacillus johnsonii* and *Parasutterella excrementihominis*, with the former being enriched in the ABX group and the latter predominating in the VH group. Functional genomic analysis of *Parasutterella excrementihominis* (Figure 5B) revealed key genomic features, including genes associated with antimicrobial resistance, metabolic pathways, and structural components, suggesting its potential role in gut dysbiosis following TBI. The genome harbored regions encoding resistance genes, rRNA, and various non-coding RNAs, emphasizing its possible contribution to the disrupted microbial landscape observed after ABX treatment. Similarly, the genomic analysis of *Lactobacillus johnsonii* (Figure 5C) identified genes linked to metabolic functions and neuroprotective properties, suggesting potential involvement in modulating host responses to TBI-induced inflammation. The presence of coding sequences for antimicrobial resistance and metabolic adaptation indicates a possible protective role of this bacterium in the ABX-treated group. These findings suggest repeated TBI induces significant gut microbial dysbiosis with severity-dependent alterations. ABX treatment modulates microbial composition and function, as evidenced by shifts in taxonomic abundance and SCFA production. These alterations suggest a potential link between gut microbiota changes and systemic inflammation in TBI pathology. Metagenomic analysis further identified the presence and enrichment of specific bacterial species, including *Lactobacillus johnsonii*, *Parasutterella excrementihominis*, and *Dubosiella newyorkensis* in the ABX-treated group, while other taxa, such as *Colidextribacter sp. OB.20* was exclusively detected in the VH-treated group (Figure S3A). Comparative genomic analysis highlights distinct microbial compositions and genomic adaptations in response to TBI and ABX interventions, underscoring the potential functional contributions of specific taxa to host-microbiome interactions post-injury. These results provide valuable insights into the gut microbiota’s role in TBI recovery and emphasize the potential for targeted microbial interventions to mitigate neuroinflammation.

**Figure 5.**
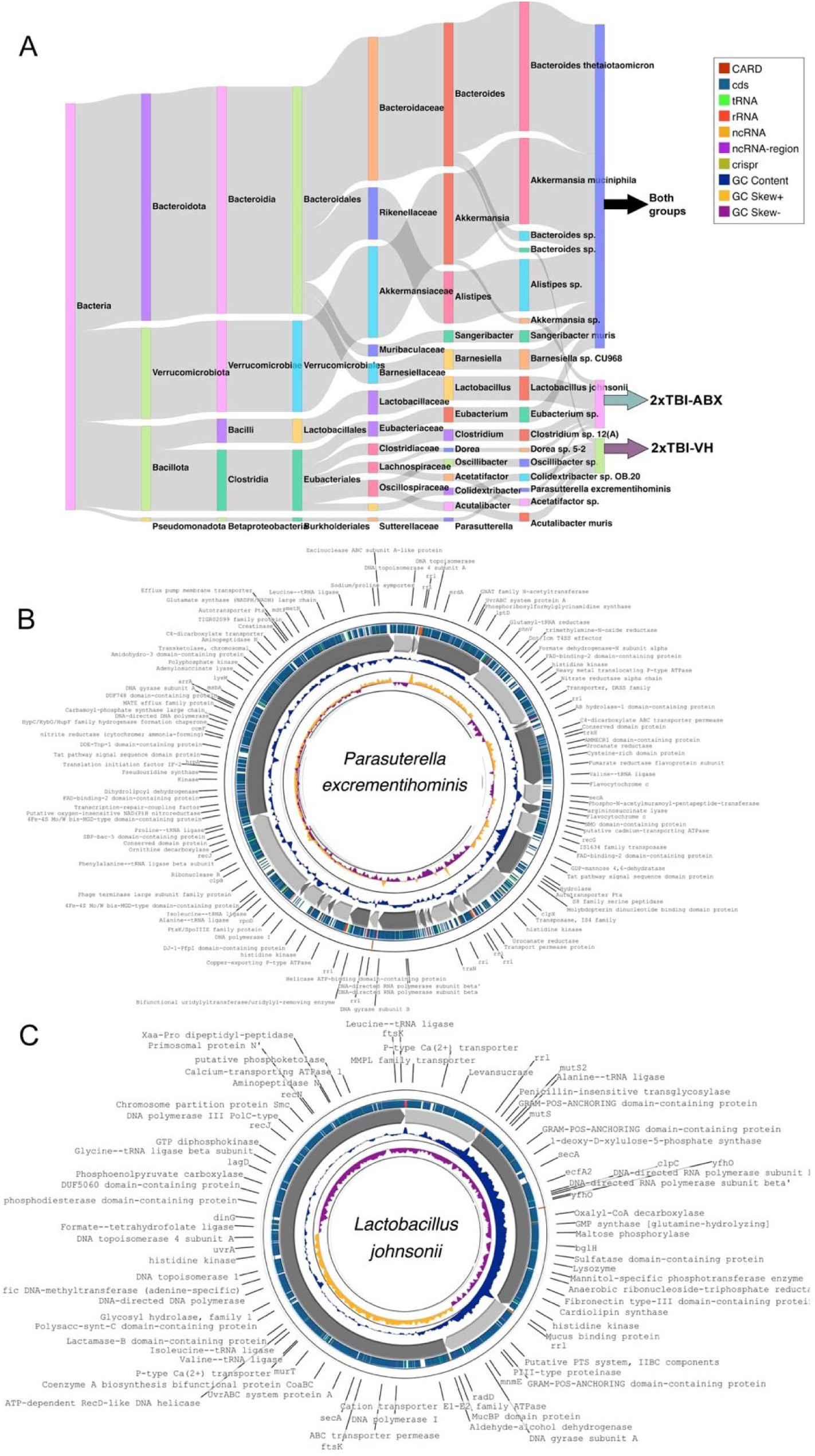
Metagenomic analysis of gut microbiome following TBI and ABX treatment. (A) Sankey diagram illustrates the taxonomic classification of bacterial communities identified in 2xTBI-ABX and 2xTBI-VH groups. n=10/group. The diagram shows the distribution of bacterial phyla, families, and genera, highlighting differences between the groups. Notably, *Lactobacillus johnsonii* and *Parasutterella excrementihominis* are more prevalent in the 2xTBI-ABX group. Both groups share common taxa, such as *Akkermansia muciniphila* and *Bacteroides* species, suggesting a core microbiome component. (B) Circular genome visualization of *Parasutterella excrementihominis*, assembled from the metagenomic analysis of the 2xTBI-ABX group. The outermost ring represents functional gene annotations, including antibiotic resistance genes (CARD, red), coding sequences (CDS, green), various RNA elements (tRNA, rRNA, ncRNA), and non-coding RNA (ncRNA). Inner rings indicate GC content, and skew illustrates genome composition (CheckM2 completeness = 97.72% and contamination = 3.68%). (C) Circular genome visualization of *Lactobacillus johnsonii*, which is more abundant in the 2xTBI-ABX group. Functional annotations include genes related to metabolic pathways, antimicrobial resistance, and neuroprotective properties. The presence of CDS, GC content, and RNA regions suggests its potential role in modulating host-microbe interactions post-TBI.

### TBI and ABX treatment induce structural alterations in gut morphology

To investigate the impact of TBI severity and ABX treatment on intestinal morphology, histological examination of ileal sections using Hematoxylin and Eosin (H&E) staining revealed notable structural differences across experimental groups. Compared to the Sh-ABX group, 1xTBI-ABX 2xTBI-ABX, and 2xTBI-VH groups displayed shortened and disorganized villi (Figure 6A). Quantification of villus morphology demonstrated a significant reduction in villi length in TBI groups compared to Sh-ABX (Figure 6B). Villi width was also significantly decreased in the 2xTBI-ABX group compared to Sh-ABX (Figure 6C), whereas crypt width was significantly decreased in the 2xTBI-ABX and 2xTBI-VH groups compared to Sh-ABX (Figure 6D). Crypt depth was mildly but significantly reduced in the 2xTBI-ABX group compared to Sh-ABX (Figure 6E).

**Figure 6.**
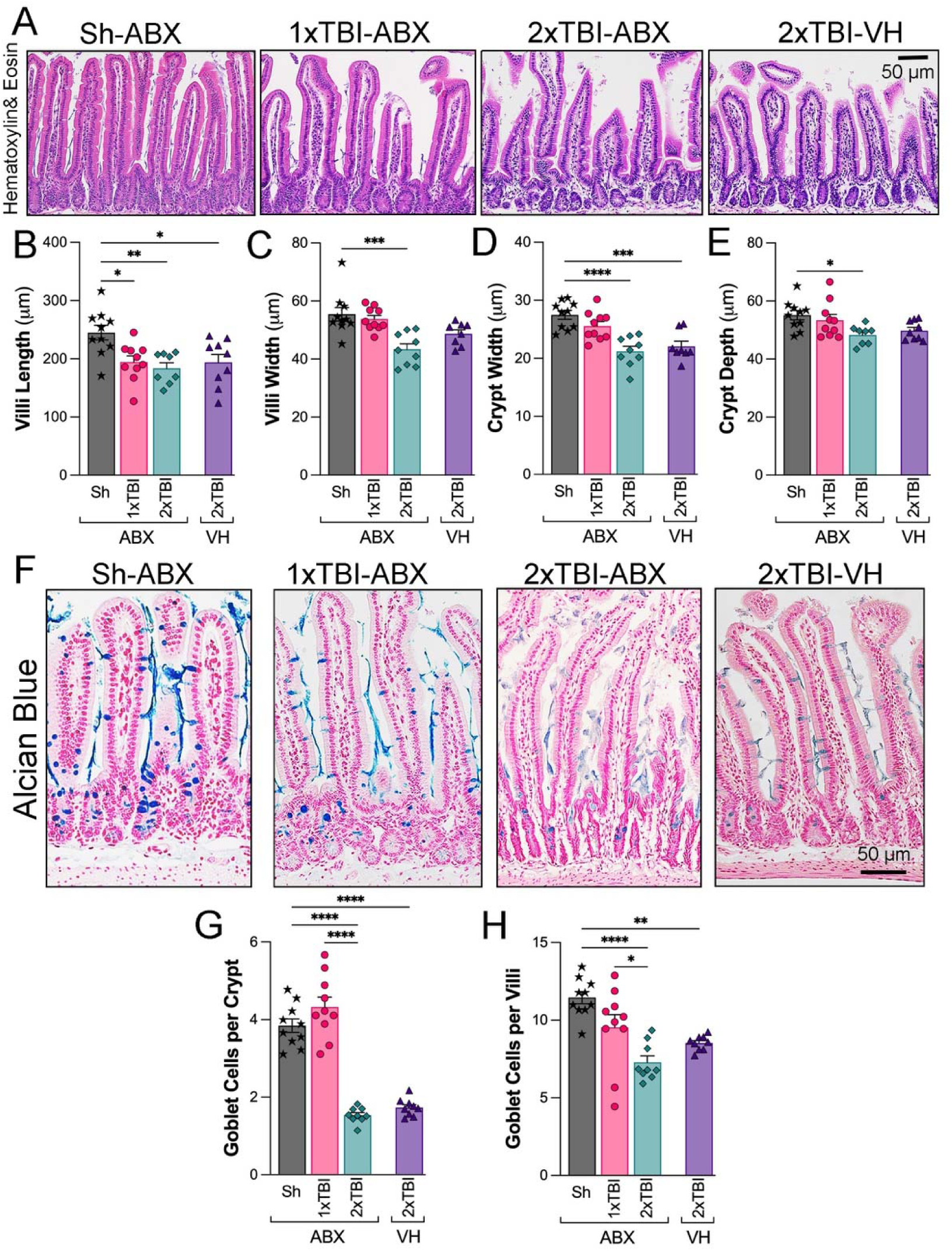
TBI and ABX treatment modulate intestinal morphology and goblet cell density. (A) Representative hematoxylin and eosin (H&E)-stained images of the small intestine across experimental groups: Sh-ABX, 1xTBI-ABX, 2xTBI-ABX, and 2xTBI-VH. (B-E) Quantification of intestinal morphology parameters: (B) Villi length (µm), showing significant shortening in the TBI groups compared to the Sh-ABX group. (C) Villi width (µm), demonstrating a reduction in the 2xTBI-ABX group compared to Sh-ABX. (D) Crypt width (µm), with a significant decrease in the 2xTBI-ABX group compared to Sh-ABX. (E) Crypt depth (µm), showing reductions in the 2xTBI-ABX compared to the Sh-ABX group. (F) Representative Alcian blue-stained images of goblet cells in the small intestine across experimental groups. Goblet cells are stained blue. (G, H) Quantification of goblet cells: (G) Goblet cells per crypt, showing a significant reduction in the 2xTBI groups compared to Sh-ABX and in the 2xTBI-ABX group compared to the 1xTBI group. (H) Goblet cells per villi significantly declined in the 2xTBI groups compared to Sh-ABX and in the 2xTBI-ABX group compared to the 1xTBI group. Data is presented as a one-way ANOVA with Tukey’s post hoc test. Data is presented as a one-way ANOVA with Tukey’s post hoc test with statistical significance indicated: *q < 0.05, **q < 0.01, ***q < 0.001, **** q<0.0001. n=10/group. Scale bar = 100 µm.

Alcian Blue staining (Figure 6F) indicated a clear reduction in goblet cell numbers and mucin production in the crypts and villi in 2xTBI groups. These cells are key components of the mucosal barrier and play important roles in host defense against intestinal pathogens. Quantification revealed a significant decrease in goblet cells per crypt (Figure 6G) and goblet cells per villi (Figure 6H) in the 2xTBI-ABX and 2xTBI-VH groups compared to both Sh-ABX and 1xTBI-ABX groups, indicating a loss of mucus-producing cells in response to repeated TBI. Together, these findings indicate that repetitive TBI has a greater effect on intestinal epithelial damage and goblet cell loss than 1xTBI or Sham groups. This may contribute to gut barrier dysfunction and dysbiosis following repeated TBI more than ABX treatment.

### Germ-free mice exhibit increased lesion volume and neuroinflammation following TBI

To investigate the impact of the gut microbiome on TBI outcomes, lesion volume, and neuroinflammatory markers were assessed in wild-type (WT) and germ-free (GF) mice. Quantification of lesion volume (Figure 7A) revealed a significant increase in GF mice compared to WT controls, indicating exacerbated tissue damage in the absence of gut microbiota. Representative DAPI-stained coronal brain sections (Figure 7B) further highlight the enlarged lesion cavity in GF mice relative to WT controls. Microglial activation was assessed using Iba-1 and CD68 immunostaining in the cortex and thalamus. Quantification of Iba-1+ cells (Figures 7C-D) demonstrated a significant increase in microglial density in the GF group compared to WT in both brain regions. Similarly, CD68+ cell counts (Figures 7E-F), a marker of activated microglia/macrophages, were significantly elevated in the cortex of the GF group, suggesting a heightened inflammatory response. Immunofluorescence images (Figures 6e1-e2, f1-f2) further support these findings, showing more significant CD68+ immunoreactivity in GF mice. GFAP staining was used to evaluate astrocytic activation following TBI. Quantification of the GFAP+ area (Figures 7G-H) revealed a significant increase in the GF group compared to WT controls, indicating enhanced astrocytic reactivity in the absence of microbiota. F4/80 staining assessed peripheral macrophage infiltration into the brain post-TBI. Quantification (Figures 7I-J) demonstrated no significant difference in macrophage infiltration in GF mice compared to WT. Immunofluorescence images (Figures 7i1-i2, j1-j2) show no differences in F4/80+ cells. P2Y12, a marker of homeostatic microglia, was significantly increased in GF mice compared to WT controls in the cortex (Figure 7K), indicating a shift toward a more activated microglial phenotype. No changes were observed in the thalamus (Figure 7L). These findings suggest that the absence of gut microbiota exacerbates lesion volume and neuroinflammation following TBI, characterized by increased microglial and astrocytic activation, heightened peripheral immune cell infiltration, and a loss of microglial homeostasis. These results highlight the critical role of the gut-brain axis in modulating the neuroinflammatory response to brain injury.

**Figure 7.**
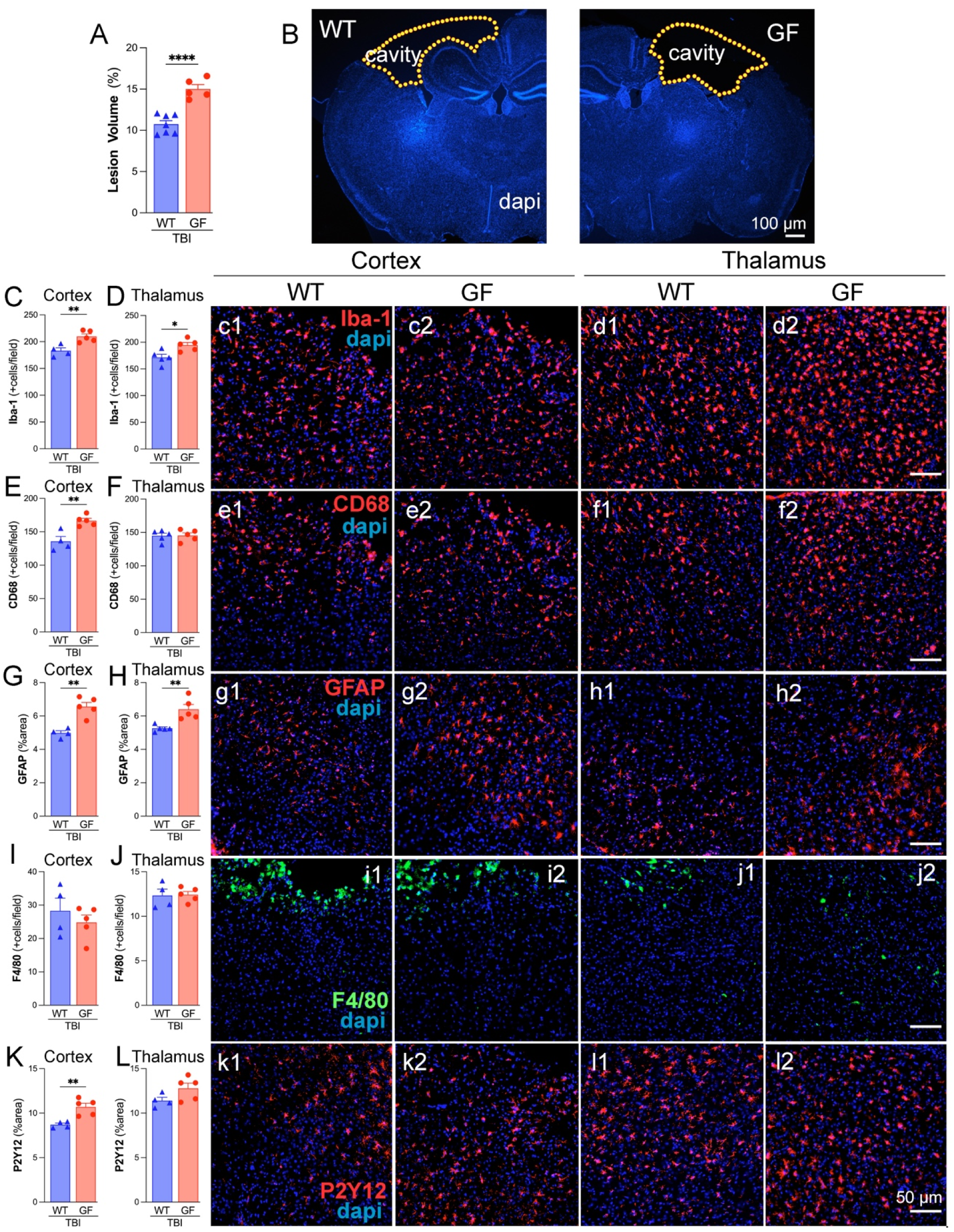
Germ-free (GF) mice exacerbate neuroinflammatory responses and lesion volume following TBI. (A) Quantification of lesion volume (%) in wild-type (WT) and germ-free (GF) mice following TBI. GF mice exhibit significantly larger lesion volumes compared to WT controls. (B) Representative images of DAPI-stained coronal brain sections of WT and GF mice showing TBI-induced lesion cavities (outlined in yellow). Scale bar = 100 µm. (C, D) Quantification of Iba-1+ microglial cells in the cortex (C) and thalamus (D), showing increased microglial activation in GF mice compared to WT controls. Representative Iba-1 (red) and DAPI (blue) images of the cortex (c1-c2) and thalamus (d1-d2). (E, F) Quantifying CD68+ microglia/macrophages in the cortex (E) and thalamus (F), demonstrating enhanced expression in the cortex of GF mice compared to WT, indicates a pro-inflammatory phenotype. Representative CD68 (red) and DAPI (blue) images of the cortex (e1-e2) and thalamus (f1-f2). (G, H) Quantification of GFAP+ astrocytes in the cortex (G) and thalamus (H) showed significant increases in GF mice compared to WT, suggesting enhanced astrocyte reactivity following TBI. Representative GFAP (red) and DAPI (blue) images of the cortex (g1-g2) and thalamus (h1-h2). (I, J) Quantification of F4/80+ macrophages in the cortex (I) and thalamus (J) showed no significant difference between the GF and WT mice. Representative F4/80 (green) and DAPI (blue) images of the cortex (i1-i2) and thalamus (j1-j2). (K, L) Quantification of P2Y12+ microglial marker in the cortex (K) and thalamus (L), showing increased expression in the cortex of GF mice, indicative of altered microglial homeostasis. Representative P2Y12 (red) and DAPI (blue) images of the cortex (k1-k2) and thalamus (l1-l2). n=4-5/group. Data is presented as a one-way ANOVA with Tukey’s post hoc test with statistical significance indicated: *p < 0.05, **p < 0.01, **** p < 0.0001. Scale bar = 100 µm (B), 50 µm (c1-l2).

## DISCUSSION

Our study demonstrates that ABX treatment reduces gut microbiome diversity but alleviates neuroinflammation following TBI. Although SCFA levels decline, the neuroprotective effects of ABX appear to occur independently of these metabolites. Additionally, we identify ABX-resilient microbial species, including *Lactobacillus johnsonii*, which may contribute to the regulation of post-TBI inflammatory responses.

ABX are commonly prescribed to TBI patients primarily to prevent infections; however, their use has been associated with the development of antibiotic-resistant infections. Clinical studies have demonstrated that factors such as ABX administration and infection significantly contribute to differences in the rectal and oral microbiomes of TBI patients compared to healthy controls ^32^. While ABX are known to compromise the immune system and overall patient health in various clinical contexts ^33–35^, their effects on brain inflammation remain largely unexplored. Our previous work showed that a single TBI event induces gut microbiome dysbiosis in mice ^20^, and that repeated TBIs sustain this dysbiotic state long-term ^36^. In the present study, we found that ABX treatment led to a marked reduction in microbiome diversity, with significant shifts observed at both the genus and species levels following repeated TBI. Interestingly, despite these microbial alterations, ABX therapy appeared to confer a net beneficial effect by attenuating neuroinflammation after TBI.

Clinical evidence indicates that TBI disrupts mucosal barrier function in ICU patients ^11^. Goblet cells, specialized epithelial cells in the gastrointestinal tract responsible for producing and secreting mucin, are essential for maintaining the integrity of the mucus barrier^37^. In our study, repeated TBI resulted in marked intestinal alterations, including goblet cell depletion, villus shortening, and blunting. Interestingly, short-term microbiome depletion appeared to exert neuroprotective effects, potentially by transiently reducing dysbiotic bacterial populations without severely compromising the gut epithelial barrier. This selective modulation of the microbiota may help dampen peripheral immune activation, thereby contributing to the observed reduction in neuroinflammatory responses.

The gut microbiota regulates BBB permeability, modulating key neurochemicals, and shaping microglial immune responses, all of which influence the brain’s reaction to injury ^38^. Studies have shown that microglia are highly responsive to gut microbiota–derived signals ^15^. In the absence of a complex microbiome, microglial maturation and function are impaired. Notably, significant genetic and morphological differences have been observed between microglia from specific pathogen-free (SPF) and GF mice ^15^. However, microbiota depletion does not enhance recovery from brain injury. In fact, GF mice, lacking a microbiome, exhibit dysfunctional microglia, an exacerbated response to acute injury, and increased tissue damage ^15^. Prior research has demonstrated that supplementation with microbiota-derived metabolites, such as SCFAs, can partially restore microglial function in GF mice ^15^. These findings underscore that a complete absence of gut microbes hinders recovery from TBI and highlight the essential role of the gut microbiome, and its metabolites, in modulating neuroinflammation and supporting brain repair after injury.

ABX decreases levels of acetate, butyrate, and propionate, which are microbial fermentation products and reduces adenine, cytosine, guanine, and uracil due to an overall reduction in bacterial load ^26^. Functional annotation of long-read metagenomic data was performed to assess SCFA level variance and its relationship with gut microbes, showing that the ABX-treated group exhibited a significant decrease in SCFA-related gene ontology (GO) term occurrences, particularly in butyrate-associated terms mainly produced by species of *Bacteroides* and *Bifidobacterium* ^39^.

In our study, we observed a reduction in circulating SCFA levels following ABX treatment. Despite this decline, neuroinflammation and brain injury were reduced, suggesting that the beneficial effects of ABX may occur through SCFA-independent mechanisms. One potential explanation is that short-term ABX treatment alters systemic immunity by depleting pro-inflammatory microbial taxa or reducing gut-derived endotoxins such as lipopolysaccharide (LPS) ^40^, thereby dampening peripheral immune activation. This reduction in systemic inflammation could subsequently mitigate neuroinflammatory responses following TBI. Interestingly, the differing outcomes between the 1xTBI+ABX and 2xTBI+ABX groups suggest the possibility of immune adaptation, tolerance, or “preconditioning” effects ^41–43^. It is plausible that the initial TBI, in the context of an altered microbiome, primes the immune system, either through the induction of tolerance or a reprogrammed inflammatory state, resulting in a distinct response to subsequent injury. This could involve mechanisms such as trained innate immunity, altered microglial activation, or modified peripheral-to-CNS signaling. Further research is needed to determine whether microbiome-driven immune memory plays a role in the divergent outcomes observed following repeated TBI.

ABX treatment modulates neuroinflammatory responses following TBI and appears to contribute to reduced brain injury severity. Previous studies have reported that administering ABX prior to neonatal hypoxia-ischemia in mice diminished early neuroinflammatory responses but did not significantly alter overall neuropathology ^44^. Consistent with these findings, our study demonstrated that ABX-treated mice showed reduced microglial/macrophage activation and decreased apoptosis, suggesting potential neuroprotective effects. However, the long-term impact of ABX treatment remains complex. Other reports have shown that while ABX may reduce lesion volume in the acute phase of TBI, it can also impair monocyte infiltration, increase neuronal loss, and elevate pro-inflammatory microglial markers months after injury ^45^. Additionally, ABX therapy disrupts gut microbiota composition by reducing overall microbial diversity, eliminating beneficial taxa, inducing metabolic shifts, increasing vulnerability to opportunistic colonization, and promoting antibiotic resistance ^46^. In this study, the selected ABX cocktail, ampicillin, gentamicin, metronidazole, and vancomycin, partially depleted the gut microbiota and led to a reduction in circulating SCFA levels following TBI. Nevertheless, certain bacterial species persisted five days after treatment, suggesting the emergence of antibiotic-resistant strains within the altered microbial community.

ABX treatment disrupts commensal gut bacteria, leading to reduced SCFA production, a decline in T helper (Th) and regulatory T (Treg) cell populations, and increased gut inflammation ^47^. Vancomycin, a narrow-spectrum ABX effective against Gram-positive bacteria like *Staphylococcus* and *Clostridium,* has been shown to diminish the production of SCFAs ^48^. In our study, we observed that both single and repeated TBI significantly reduced microbiota diversity and richness, with the most dramatic changes occurring within the first 24 hours post-injury, consistent with previous findings in TBI rodent models ^49^.

Following a single TBI, we observed a significant decrease in beneficial bacteria such as *Bifidobacterium* and *Lactobacillus* ^50^. With repeated TBI, this trend continued and extended to other genera, including *Clostridium* and *Turicibacter*, paralleling an increase in neuroinflammatory responses. These taxa are notable for their roles in gut-brain communication; for example, *Lactobacillus* species have been shown to modulate vagus nerve activity, influence levels of neurotransmitters such as GABA, glutamate, and serotonin, and reduce anxiety-like behaviors in mice ^51^. Similarly, both *Bifidobacterium* and *Lactobacillus* are known GABA producers ^52^, suggesting their potential neuroprotective roles post-TBI—a finding supported by our earlier studies ^53^. Another study suggested that *Lactobacillus* may play a neuroprotective role by reshaping the intestinal microbiota in TBI mice ^54^. These findings indicate that TBI disrupts the balance between beneficial and harmful bacteria, which may significantly impact brain recovery. Interestingly, ABX administration led to a relative enrichment of *Akkermansia* in both control and single TBI groups. As a mucin-degrading microbe, *Akkermansia* is increasingly recognized for its role in gut-brain axis signaling ^55^.

In both single and repeated TBI groups treated with ABX, we also observed an increase in *Parasutterella* and a decrease in *Dubosiella*. Additionally, *Bacteroides* levels rose in both vehicle and ABX-treated groups, possibly reflecting a compensatory role in post-TBI repair via its known capacity to produce GABA ^56^. Through 16S rRNA sequencing, *Parasutterella* emerged as a notable outlier in ABX-treated animals. Metagenomic reconstruction identified the species as *Parasutterella excrementihominis*, for which we successfully assembled a high-quality metagenome-assembled genome (MAG) (Figure 5B). Gene annotation suggests that this species may exert anti-inflammatory effects, consistent with previous studies showing its role in reducing adipose tissue inflammation in mice ^57^. We also detected *Lactobacillus johnsonii* (Figure 5C) in the ABX group at 5 dpi. Our prior work demonstrated that fecal microbiota transplants (FMT) from Alzheimer’s model mice into healthy controls reduced *L. johnsonii* abundance and heightened microglial activation after TBI ^58^. Notably, *Lactobacillus* species ^59^, including *L. johnsonii*, exhibit high rates of antibiotic resistance ^60^. Clinically, *L. johnsonii* has been proposed as a promising immunomodulator and potential therapeutic target for conditions like ulcerative colitis ^61^. While we did not observe increased SCFA levels, *L. johnsonii* likely contributes to reduced peripheral inflammation by promoting Treg responses, inhibiting pro-inflammatory Th-cell activation, and stimulating anti-inflammatory cytokines like TGF-β ^62^.

Moreover, administration of *L. johnsonii* BS15 has been shown to improve neuroinflammation and demyelination in the hippocampus and to alleviate memory dysfunction in murine models of fluoride exposure ^63^. Extracellular vesicles secreted by *L. johnsonii* have also demonstrated anti-inflammatory effects by modulating macrophage phenotypes in response to pathogenic *E. coli* strains ^64^. Given its anti-inflammatory properties and inherent antibiotic resistance, *L. johnsonii* represents a promising candidate for modulating post-TBI neuroinflammation. Future studies should investigate the therapeutic potential of *L. johnsonii* supplementation following TBI.

### Limitations of the study

This study was conducted using only male mice, which limits generalizability and does not account for potential sex-specific effects. Cage effects and the nature of the behavioral assays, some of which may influence gait, should also be considered. The use of short-read 16S rRNA amplicon sequencing restricted taxonomic resolution to the genus level ^65^, although this was mitigated by complementary long-read metagenomic sequencing ^66^, which enabled species- and strain-level identification. Nevertheless, functional predictions remain limited by sequencing depth and by the complex regulation of microbial metabolite production, particularly SCFAs, which are central to gut-brain axis communication.

### Conclusions

TBI induces gut dysbiosis in a severity-dependent manner, altering microbial populations in ways that may exacerbate systemic and neuroinflammation. Short-term ABX treatment can attenuate these microbial shifts and reduce neuroinflammatory responses, albeit through SCFA-independent mechanisms. Our findings highlight the potential anti-inflammatory roles of *L. johnsonii* and *Parasutterella excrementihominis* as modulators of post-TBI inflammation. Probiotic interventions targeting these species represent promising therapeutic avenues for improving recovery after TBI.

## Ethics approval and consent to participate

The Animal Care and Use Committee of Houston Methodist Research Institute, Houston, TX, USA, approved all animal experiments in the study.

## Consent for publication

Not applicable.

## Author Contributions

H.F., A.M., M.H., M.B., L.C., S.S., T.J.T., and S.V. initiated, designed, planned, and oversaw all study aspects. H.F., A.M., M.H., M.B., L.C., and S.V. performed the experimental work and data analysis, and H.F. and S.V. drafted the manuscript. All authors reviewed and edited the final version of the manuscript.

## Acknowledgments

This study was supported by NIH grant R21NS106640 (S.V.) from the National Institute for Neurological Disorders and Stroke (NINDS) and NIH grant R56AG080920 (S.V.) from the National Institute on Aging (NIA). The content is solely the responsibility of the authors and does not necessarily represent the official views of the NIH. A.M. is supported by a training fellowship from the Gulf Coast Consortia on the NLM Training Program in Biomedical Informatics & Data Science (T15LM007093). The authors thank the CPRIT Proteomics and Metabolomics Core Facility (RP210227), NIH (P30 CA125123), and Dan L. Duncan Cancer Center at Baylor College of Medicine. We also thank the Pathology and Histology Core at Houston Methodist Research Institute. We also thank Dr. Paula Scalzo for her valuable assistance in reviewing the statistical analyses presented in this study. Figure 1a was created using Biorender.

## STAR⍰Methods

### Key resources table

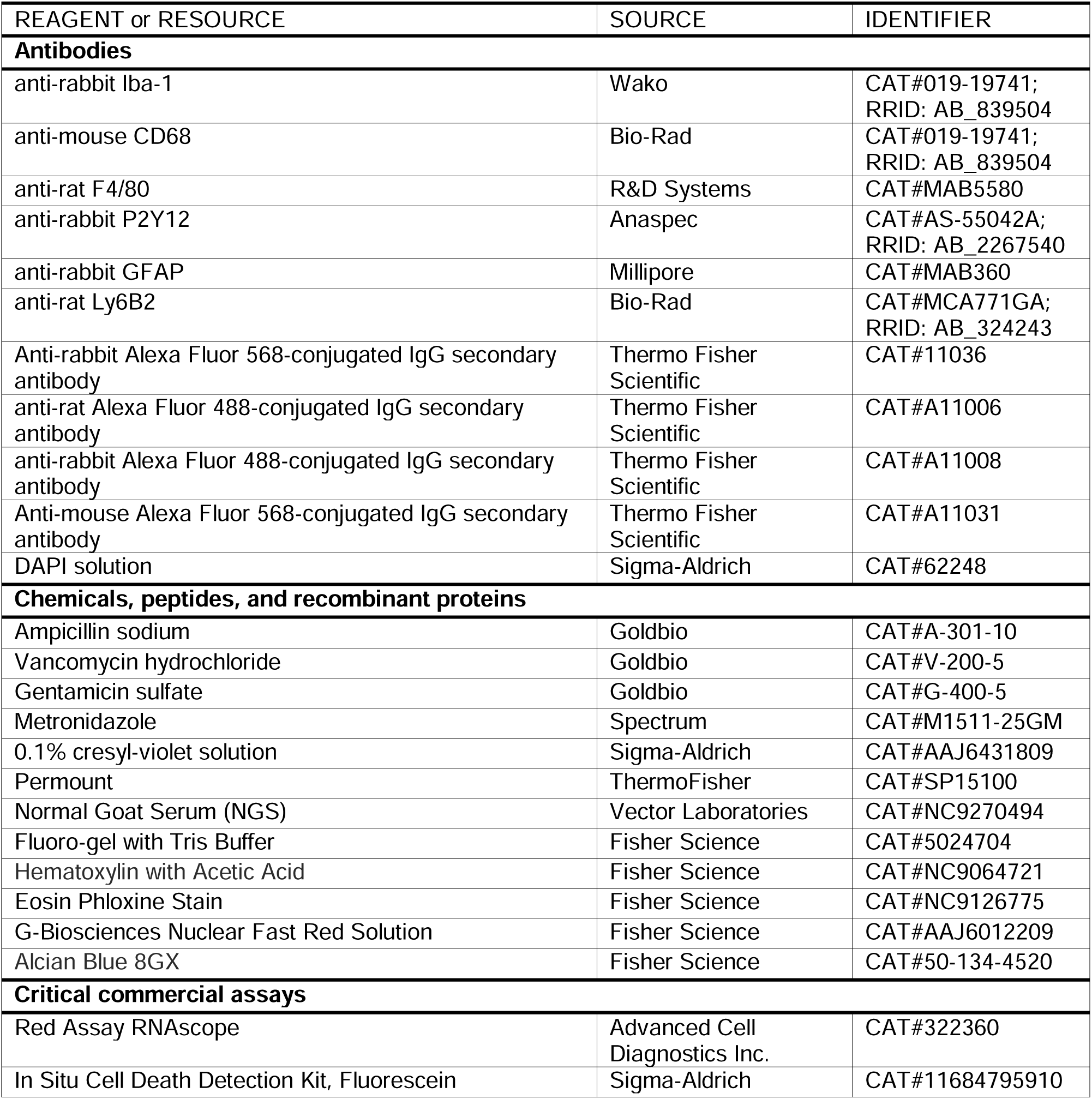
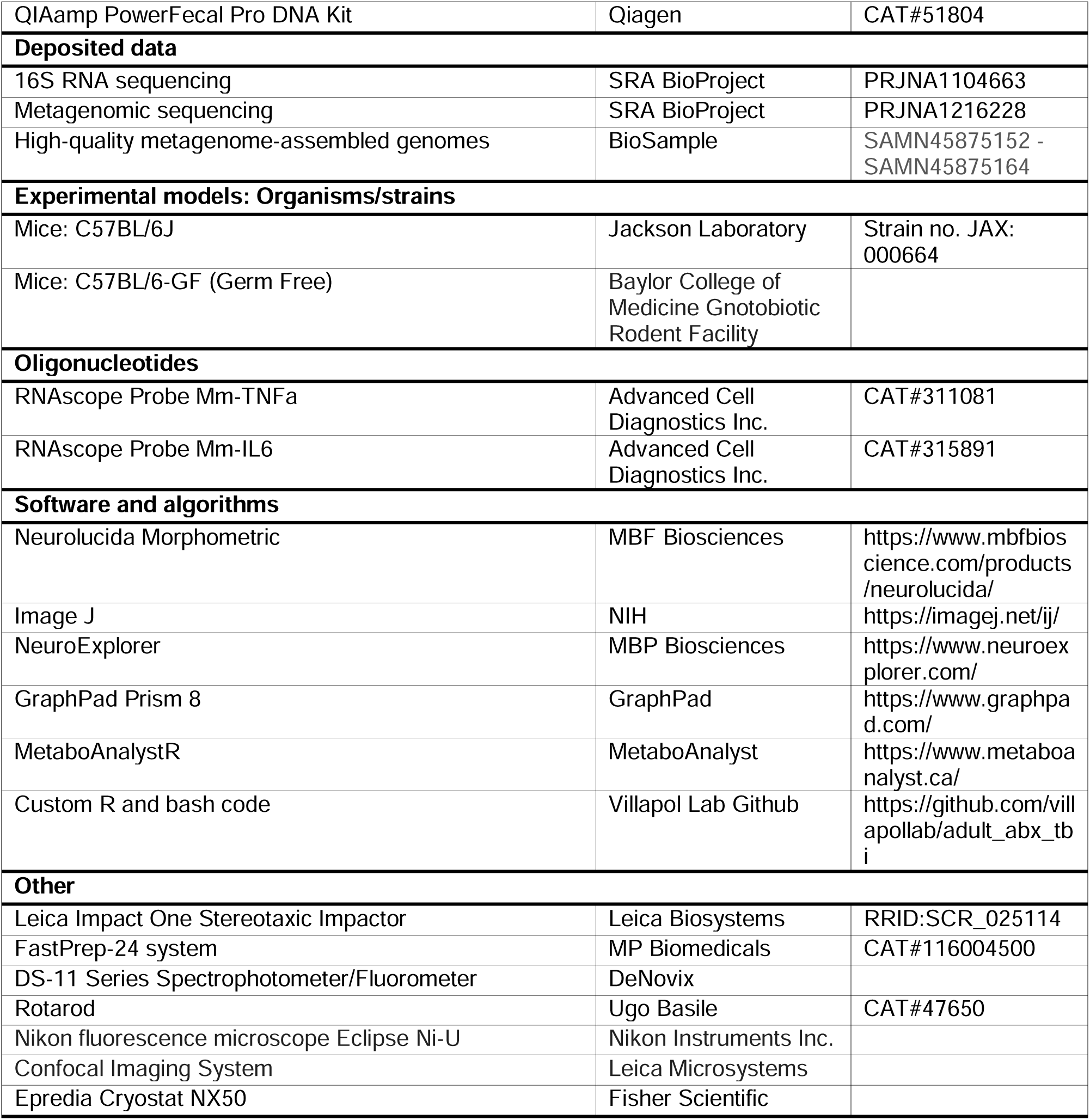

### Resource Availability

#### Lead Contact

Further information and requests for resources and reagents should be directed to and will be fulfilled by the lead contact, Dr. Villapol.

#### Materials availability

- All materials generated in this study are available from the lead contact upon request:

#### Data and code availability

- The 16S RNA sequencing and metagenomic profiling datasets generated in this study are in the NCBI SRA database, BioProject numbers PRJNA1104663 (raw 16S sequencing data and MAG assemblies) and BioProject PRJNA1216228 (raw metagenomic sequencing data).
- The code used for the amplicon and metagenomic analysis can be found in the document on our GitHub page: https://villapollab.github.io/adult_abx_tbi/.
- The lead contact can provide any additional information required to reanalyze the data reported in this paper upon request.

### Experimental model and study participant details

#### Mice

Young adult (14-week-old) wild-type C57BL/6J male mice (Jackson Laboratories, Bar Harbor, ME) were housed at the Houston Methodist Research Institute animal facilities under a standard 12-hour light and dark cycle with access to food and water ad libitum. All *in vivo* experiments were approved by the Institutional Animal Care and Use Committee (IACUC) at Houston Methodist Research Institute, Houston (TX, USA). After arriving, the mice were given one week of acclimatization to adjust to their new surroundings before participating in any experimental procedures. Adult male C57BL/6 mice were divided into groups for the study. We included only male mice due to findings indicating males exhibit a higher inflammatory response after TBI than females ^67^. Germ-free (GF) mice strains were used to prove inflammatory comparison between the ABX-induced microbiome depletion mice. C57BL/6-GF male mice (12 weeks old) were obtained from the Baylor College of Medicine Gnotobiotic Rodent Facility (Houston, TX). An internal standard was serially diluted to assess the “*germ-free*” status of the mice upon arrival, and the copy number of the 16S rRNA gene in feces from each transfer crate was analyzed using quantitative PCR (qPCR). 16S rRNA gene was not detected in feces from transfer crates, confirming that the mice were GF (data not shown).

### Method details

#### ABX treatment

To achieve microbiome depletion in the C57BL/6 mice, an ABX cocktail was made consisting of ampicillin (1 mg/mL), gentamicin (1 mg/mL), metronidazole (1 mg/mL), and vancomycin (0.5 mg/mL), dissolved (key resources table) in autoclaved drinking water. The cocktail was administered to ABX-treated male mice via oral gavage at 200 µL for 3 consecutive days. The vehicle (VH)-treated mice underwent all procedures except microbiota depletion and were administered water as control VH via oral gavage.

#### Traumatic brain injury model

Mice from both TBI groups were anesthetized with isoflurane before receiving an initial controlled cortical impact (CCI) injury on the left hemisphere at the primary motor and somatosensory cortex using an electromagnetic Impact One stereotaxic impactor (Leica Biosystems, Buffalo Grove, IL, USA). The impact site was localized at 2 mm lateral and 2 mm posterior to Bregma with a 3 mm diameter flat impact tip. The impact was made at a velocity of 3.2 m/s and impact depth of 1.5 mm. As specified in our earlier studies, these parameters were determined to induce an inflammatory response post-TBI ^2,67^. Sham mice were undergoing preparation for TBI but did not receive impacts. The inclusion criteria for TBI mice involved confirmation of injury by righting reflex time (>5 min) and post-injury behavioral responsiveness. Mice that did not meet these criteria or experienced complications during the procedure were excluded. We established two experimental models to investigate whether a healthy mouse microbiome responds differently to acute TBI than a dysbiotic microbiome from an injured animal. One group received a single CCI (1xTBI), while another underwent a second CCI 38 days after the initial injury (2xTBI) (Figure 1A). These models were designed to minimize the impact of injury severity on microbiome composition, inflammatory responses, and the subsequent effects of ABX treatment in previously injured animals. The experimental groups receiving ABX treatment included Sh, 1xTBI, and 2xTBI mice.

Additionally, to assess the influence of ABX on recovery, we incorporated a VH-treated control group with 2xTBI. Mice were anesthetized and euthanized at 5 days post-injury (dpi) and ABX treatment. We exposed GF mice and wild-type (WT) mice to CCI injury using the same parameters, and then the mice were euthanized at 3 dpi to assess the impact of acute TBI on neuroinflammation and lesions. Brains, blood, and small intestines were collected for further analysis.

#### Rotarod test

Sensorimotor function was evaluated using a Rotarod behavior test from Ugo Basile (Gemonio, Italy) to evaluate the impact of ABX on motor function. As previously described, mice underwent an initial training session consisting of three trial runs conducted two days before behavior testing. During each trial, individual mice were placed on the rotarod apparatus and trained until they could remain on the rod for a minimum of 30 seconds without falling. To standardize performance assessment, a maximum latency of 300 seconds was imposed. Mice displaying passive rotation behavior, clinging to the rod without active ambulation, were excluded from analysis, as this behavior does not represent coordinated motor function ^68–70^. Mice were then assessed 1 and 5 days after ABX or VH treatment.

#### Fecal Microbiome DNA extraction for 16S rRNA Sequencing

Gut microbiome DNA concentrations were measured through DNA extraction from fresh frozen stool samples from the VH and ABX groups at the following time points: -38 days, -3 days, 0 days, and 5 days post-treatment (Figure 1A). After collection, stool samples were stored at −80°C. Genomic bacterial DNA was extracted from frozen stool samples using the QIAamp PowerFecal Pro DNA Kit (Qiagen, Germantown, MD). Bead beating was implemented in three cycles for DNA extraction, each lasting one minute at a speed of 6.5 m/s and a rest period of five minutes between cycles, using a FastPrep-24 system (MP Biomedicals, Irvine, CA). DNA isolation was completed according to the manufacturer’s instructions. The concentration of the extracted genomic DNA was then measured using a DS-11 Series Spectrophotometer/Fluorometer (DeNovix, Wilmington, DE). If bacterial DNA extraction from fecal material yielded undetectable levels, these samples were not included in downstream quantification. Extracted genomic bacterial DNA from collected fecal samples was analyzed for microbiota colonization and diversity by 16S rRNA gene compositional analysis. Illumina MiSeq was performed using the adapters and single-index barcodes so that the polymerase chain reaction (PCR) products could be pooled and sequenced directly, targeting at least 10,000 reads per sample ^71^. Primers used for the 16S V1-V3 amplification were 27F (AGAGTTTGATYMTGGCTCAG, where Y = C (90%) or T (10%); M = A (30%) or C (70%) and 534R (ATTACCGCGGCKGCTGG, where K = G (10%) or T (90%) ^72^. Amplicons were generated using primers corresponding to the V1-V3 variable regions, and the PCR products were purified. Subsequently, sequencing libraries for the V1-V3 target were constructed following the instructions provided by the Illumina MiSeq system with end products of 300 bp paired-end libraries.

#### 16S microbiome data analysis

Raw data files in binary base call (BCL) format were converted into FASTQs and demultiplexed based on the single-index barcodes using the Illumina ‘bcl2fastq’ software. Demultiplexed read pairs underwent quality filtering using bbduk.sh (BBMap version 38.82), removing Illumina adapters, PhiX reads, and sequences with a Phred quality score below 15 and length below 100 bp after trimming. 16S V1-V3 quality-controlled reads were then merged using bbmerge.sh (BBMap version 38.82), with merge parameters optimized for the 16S V1-V3 amplicon type (vstrict=t qtrim=t trimq=15). Further processing was performed using custom R and bash scripts, found in our documentation at https://villapollab.github.io/adult_abx_tbi/. Sequences were processed sample-wise (independent) with DADA2 v1.32 to eliminate any residual PhiX contamination, trim reads (forward reads at 275 bp and reverse reads at 265 bp; reads shorter than this were discarded), discard reads with > 2 expected errors, correct errors, merge read pairs, and to remove PCR chimeras. After clustering, 12,008 amplicon sequencing variants (ASVs) were obtained across all samples. The ASV count table contained a total of 2,589,891 counts, at least 4,519 and at most 42,595 per sample (average 18,767). Taxonomic classification was performed in DADA2 with the assigned taxonomy function using the precompiled GreenGenes2 release 2024.09 databases ^73^. The resulting ASV matrix, taxonomy table, and metadata table were converted to a phyloseq v1.48.0 object and merged with the phylogenetic tree externally calculated from the ASVs using MAFFT v7.525 and FastTree v2.1.11. The phyloseq object was used in the creation of alpha and beta diversity plots, PERMANOVA calculations with vegan::adonis2 v2.6-5, relative abundance bar plots with microViz v0.12.5, R v4.3.3, and differentially abundant taxa calculations using ANCOMBC2 v2.6.0.

#### Shotgun metagenomic sequencing and data analysis

Nanopore long-read metagenomic sequencing (R9.4.1 chemistry) was performed on two pools of mice fecal DNA from the 2xTBI-ABX and 2xTBI-VH groups at 5 dpi. The samples were selected from mice within the same cage for each treatment group. The SQK-RAD004 library preparation kit was used with equal volumes from each mouse taken to sum to ∼8 μL of pooled fecal DNA, which was processed following the recommended protocol and loaded into each flow cell (∼1 μg per flow cell). Raw nanopore sequencing data were output in pod5 format and basecalled using Dorado v0.7.0 with the super accuracy model. After converting to fastq format with samtools v1.21, quality and length filtering and host-read removal were performed using NanoPlot v1.43.0, hostile v1.1.0, and chopper v0.9.0 ^74–76^. Once fastq files were thoroughly cleaned and trimmed, metagenomic assembly, read mapping, binning, and genomic taxonomic classification were performed using the snakemake pipeline Aviary v0.10.0 ^77–83^. Functional annotation of the fastq files for each treatment group was performed using SeqScreen v4.4 with a custom R script to parse the functional_results.txt output and compare treatment groups on a list of relevant GO terms ^84^. The combination of lemur and magnet was used to verify if a bacterial taxon was truly present when not able to construct a draft metagenomic bin ^85^. Metagenome-assembled genomes (MAGs) were generated for 20 of the most abundant taxa across all samples using the MIMAG recommendations of CheckM2 Completeness ≥ 90% and Contamination ≤ 5% for high-quality MAGs and draft quality MAGs having scores of Completeness ≥ 80% and Contamination ≤ 10%.

#### Serum SCFA analysis

SCFAs were analyzed by derivatization from collected blood serum samples from 2xTBI-ABX and 2xTBI-VH groups at 5 dpi. Blood serum was isolated via centrifugation (15 min x 3,000g). Once extracted, 40 µL of serum was added to 40 µL of acetonitrile, vortexed, and centrifuged. 40 µL of the supernatant, 20 µL of 200 mM 12C6-3-Nitrophenylhydrazine (3NPH), and 120 mM 1-Ethyl-3-(3-dimethylaminopropyl)carbodiimide (EDC) were combined. 20 µL of hydrochloric acid was added and incubated for 30 min at 40°C. The resulting mixture was cooled and made up to 1.91 mL with 10% aqueous acetonitrile. 5 µL of the sample was injected into liquid chromatography-tandem mass spectrometry (LC-MS/MS). SCFAs were separated using mobile phases: 0.1% formic acid in water (mobile phase A) and 0.1% formic acid in acetonitrile (mobile phase B). Separation of metabolites was performed on Acquity UPLC HSS T3 1.8 um (2.1×100mM). The SCFA profiles were measured in ESI negative mode using a 6495 triple quadrupole mass spectrometer (Agilent Technologies, Santa Clara, CA) coupled to an HPLC system (Agilent Technologies, Santa Clara, CA) with multiple reaction monitoring (MRM). The acquired data was analyzed using Agilent Mass Hunter quantitative software (Agilent Technologies, Santa Clara, CA). Raw peak intensity for each SCFA was normalized by sum, log transformed, and auto-scaled (mean centered and divided by standard deviation).

#### Lesion volume and cell death assay

To evaluate the severity of injury and the effects of single or repeated TBIs, as well as the influence of ABX treatment, lesion volume was quantified in the ipsilateral hemisphere. Whole brain samples were fixed and sectioned using a cryostat (Epredia Cryostar NX50, Fisher Scientific, Waltham, MA, USA). Coronal sections, 16 μm thick, were either mounted directly onto glass slides or immersed in a free-floating cryoprotective solution containing 30% sucrose, 1% polyvinylpyrrolidone, 30% ethylene glycol, and 0.01M PBS. For immunohistochemical usage, collected sections were taken at coronal planes from the frontal cortex throughout the dorsal hippocampus. A cresyl-violet solution was prepared under a ventilated hood by combining 0.1% cresyl-violet (Sigma-Aldrich, St. Louis, MO, US), acetic acid, and distilled water. Pre-mounted brain sections were stained with cresyl violet solution for 10-20 min before dehydrating in ethanol dilutes. The slides were submerged in xylene before being covered with a xylene-based Permount (ThermoFisher Scientific) mounting media and coverslipped. The lesion volume was calculated as a percentage of the lesion area. Collected data was then averaged for each of the 9-12 brain sections per slide using ImageJ software. Cell death was assessed using the Fluorescence *In Situ* Cell Death Detection Kit (Sigma-Aldrich, Indianapolis, IN, USA). DNA strand breakage in brain sections was analyzed via Terminal deoxynucleotidyl transferase dUTP nick end labeling (TUNEL), following the manufacturer’s protocol ^2^. During the TUNEL staining process, some brain sections were compromised due to brain tissue drying or loss during slide mounting. These issues affected tissue integrity and prevented reliable quantification, so the corresponding brains were excluded from analysis. To ensure data quality and consistency, 1 to 3 animals were discarded per group based on these criteria. Histological images were captured using a Nikon fluorescence microscope (Eclipse Ni-U, Melville, NY, USA) and a confocal imaging system (Leica Microsystems, Deerfield, IL, USA). The regions of interest included the somatosensory cortex and thalamus.

#### Immunofluorescence staining and *in situ* hybridization

Free-floating brain sections were first washed with a series of PBS and 0.5% PBS-Triton X-100 (PBS-T) before applying a 3% normal goat serum (NGS) (Vector Laboratories, Burlingame, CA) blocking solution for 1 hour at room temperature (RT). A primary antibody solution made from blocking solution (PBS-T and 3% NGS) and dilutions of the following primary antibodies: anti-rabbit Iba-1 (1:500, Wako), anti-mouse CD68 (1:200, Biorad), anti-rat F4/80 (1:200, R&D Systems), anti-rabbit P2Y12 (1:500, Anaspec), anti-rabbit GFAP (1:500, Millipore), and anti-rat Ly6B2 (1:500, Biorad), were incubated at 4 °C overnight. The next day, brain sections were washed in PBS-T 3×5min and incubated with the corresponding anti-rabbit or anti-mouse Alexa Fluor 568-conjugated and anti-rat or anti-rabbit Alexa Fluor 488-conjugated IgG secondary antibody (1:1000, Thermo Fisher Scientific, Waltham, MA, USA) for 2 hours at RT. Samples were washed with distilled water 3×5min before being counterstained with DAPI solution diluted in PBS (1:50,000, Sigma-Aldrich) for 5 min. Sections were mounted, and the cover slipped using Tris Buffer mounting medium (Electron Microscopy Sections, Hatfield, PA). Fluorescent *in situ* hybridization (FISH) was performed as per the manufacturer’s instructions using RNAscope® Technology 2.0 Red Fluorescent kit (Advanced Cell Diagnostics (ACD), Hayward, CA, US) as previously described ^67,86^. Brain tissue sections were dehydrated using an ethanol series of 50%, 70%, and two times 100% for 5 min. Subsequently, they were boiled for 10 minutes with pretreatment 2 solution (citrate buffer). The slides were incubated with pretreatment 3 solution (protease buffer) for 30 min before hybridization. For hybridization, sections were incubated at 40 °C for 2 h with specific target probes: Mus musculus TNF-α (Cat. No.311081, ACD) and Mus musculus il-6 (Cat. No. 315891. ACD). In addition, the negative (Cat. No. 310043, ACD) and positive (Cat. No. 313911, ACD) control probes were applied and allowed to hybridize for 2 h at 40 °C. After FISH, the slides were co-stained with anti-rabbit Iba-1 as previously described^87^.

#### Quantitative analysis

We employed unbiased, standardized sampling methods to evaluate tissue areas in the cortex and thalamus that exhibited positive immunoreactivity for the quantitative analysis of immunolabeled sections. To measure the number of Iba-1+ cells, we analyzed an average of five single-plane sections from the lesion center (ranging from −1.34 to −2.30 mm from bregma) for each animal, blind to the conditions across each brain region. Within each region, all cells positive for Iba-1, CD68, F4/80, and Ly6B2 were counted in five specific fields in the cortex and two specific fields in the thalamus (x20, 151.894 mm²) near the impact site. For proportional area measurements, we quantified the extent of the reaction for microglial and astroglia cells as the percentage of the area occupied by immunohistochemically stained cellular profiles within the injured cortex and thalamus regions. The data were presented as the percentage of the area showing P2Y12 or GFAP+ immunoreactivity relative to the total studied area across 15 cortical and thalamus areas at the impact level. The quantitative image was performed using ImageJ64 software (NIH, Bethesda, MD, USA) as previously described ^67^ for inversion, thresholding, and densitometric analysis. The threshold function was utilized to set a black-and-white threshold corresponding to the imaged field, subtracting the averaged background. The “Analyze Particles” function was then used to calculate the total area of positive staining and the proportion of the total area.

#### Assessment of microglia morphological analysis

Immuno-stained Iba-1 brain sections were imaged at a 40x oil objective using a confocal imaging system (Leica Microsystems, Deerfield, IL, USA) to further investigate microglia structure and morphology. Microglia in the somatosensory cortex and thalamus were reconstructed with Neurolucida morphometric software (MBF Biosciences, VT, USA) by using a 3D sectional plane. First, microglia somas were individually labeled to identify the central points of each microglia. Dendrite branch mapping was then performed using the program’s tracing tool to label dendrite thickness, curvature, and density. Once mapped, tracings were rendered into a 2D diagram using NeuroExplorer software (MBP Biosciences, VT, USA). A Sholl analysis assessed cell complexity concerning soma size and distance. Concentric circles spaced 2 μm apart, originating from the soma, were placed over each microglia. The number of dendrite branches that intersected the radius, the average dendrite length, the number of nodes (branching points), and the average surface area of individual microglia were measured as a function of the distance from the cell soma for each radius, giving an idea of glial activity and function (Supplementary Figure 1).

#### Assessment of gut histology

The small intestines of grouped mice were isolated and fixed in 4% paraformaldehyde for 24 hours, followed by dehydration in a 70% ethanol solution. Tissue samples were processed using a Shandon Excelsior ES Tissue Processor and embedded in paraffin with a Shandon HistoCenter Embedding System, following the manufacturer’s standard protocols. The samples were sectioned at a thickness of 5 μm and mounted onto glass slides. Hematoxylin and Eosin (H&E) staining was performed to assess tissue structure. Intestinal sections were deparaffinized in xylene, rehydrated in water, and stained with hematoxylin for 6 hours at 60–70°C. After rinsing with tap water to remove excess stain, the sections were differentiated using 0.3% acid alcohol for 2 minutes, followed by eosin staining for 2 minutes. The slides were then rinsed and mounted with a xylene-based Permount mounting medium, allowing them to dry overnight. Alcian Blue staining was conducted to evaluate mucin production by goblet cells in the intestines. Deparaffinized sections were dehydrated in graded ethanol solutions and washed in distilled water before applying the Alcian Blue solution for 30 minutes. Excess stain was removed with tap water, followed by counterstaining with Nuclear Fast Red Solution for 5 minutes. Samples were then rinsed, dehydrated, cleared in xylene, and mounted as previously described ^58^.

### Quantification and statistical analysis

#### Statistical analysis

The sample size chosen for our animal experiments in this study was estimated based on our prior experience performing similar experiments. Statistical analysis was performed using unpaired Student’s t-tests for two groups (GF vs. WT groups), and one-way analysis of variance (ANOVA) or two-way ANOVA for multiple groups. Partial least squares discrimination analysis (oPLS-DA) calculation and visualization were performed using the MetaboAnalystR [56,58] package within R (version 4.3.3). For the SCFAs, the distribution was non-parametric, so we used the Mann-Whitney test, which assesses data distribution in ranks. Analysis of the rotarod test and histochemical/immunofluorescence utilized a two-way ANOVA followed by Tukey’s multiple comparisons to compare the time after injury and sex as the independent variables. All mice were randomly assigned to experimental conditions, and experimenters were blinded to the treatment groups throughout the study. Statistical analyses were conducted using GraphPad Prism 8 software (GraphPad, San Diego, CA, USA) for multiple groups, with all data points showing a normal distribution. Data are presented as significance levels set at *p < 0.05, **p < 0.01, ***p < 0.001, and ****p < 0.0001. The sample sizes and the main effects of the statistical details of experiments can be found in each figure legend.

---

**Supplementary Figure 1.**
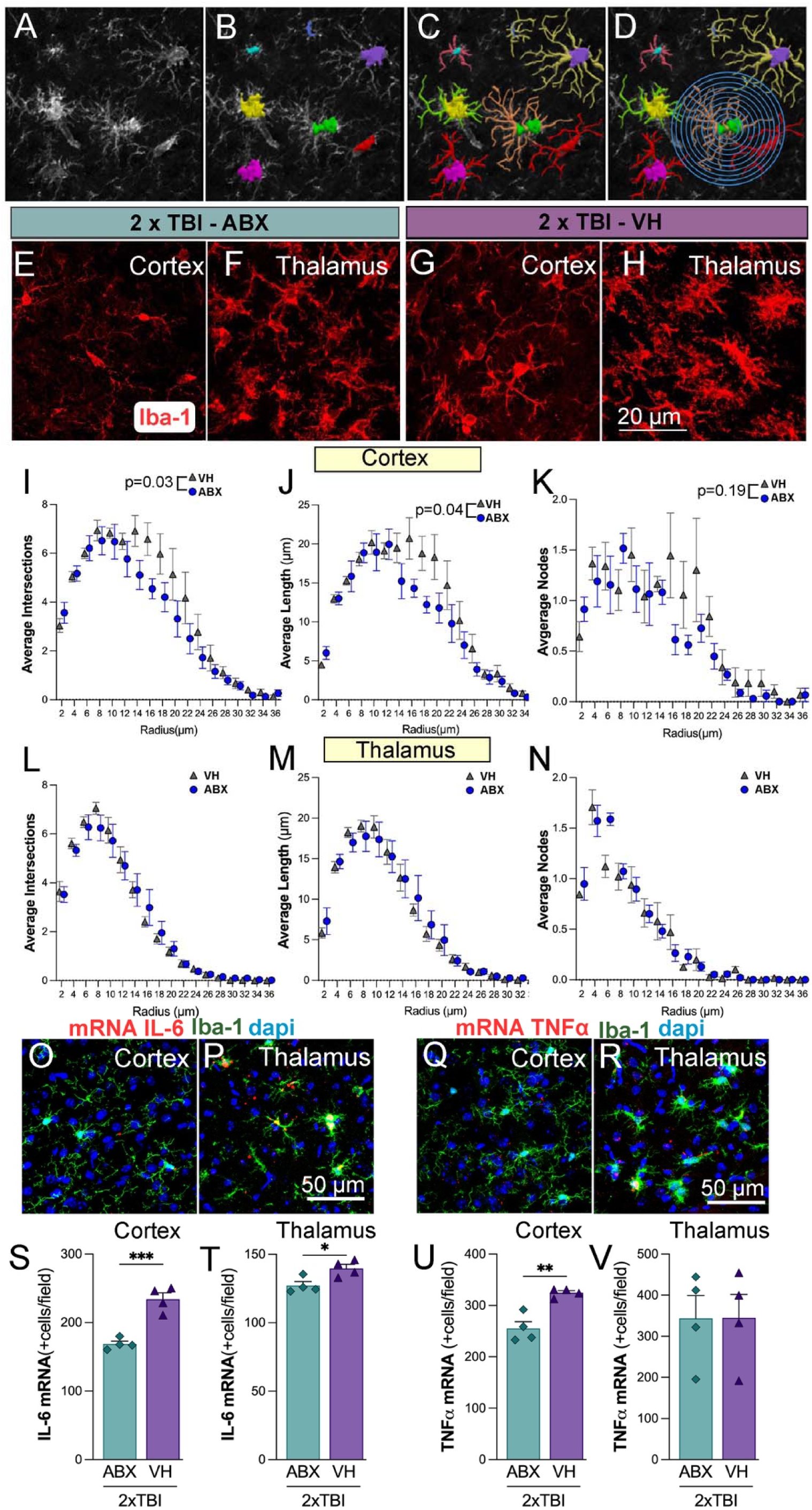
Microglial morphological changes following repeated TBI with and without ABX treatment. (A-D) Representative images of microglial morphology analysis in the 2xTBI-VH group. (A) Original grayscale Iba-1 staining image, (B) segmented microglia with color-coded individual cells, (C) skeletonized microglia used for Sholl analysis, and (D) overlay of Sholl analysis with concentric circles to assess microglial branching complexity. (E-H) Representative Iba-1 immunofluorescence images of microglia in the cortex (E, G) and thalamus (F, H) from 2xTBI-ABX and 2xTBI-VH groups, respectively. Increased microglial activation is observed in the VH-treated group compared to the ABX-treated group. (I-N) Sholl analysis quantifying microglial morphology in the cortex (I-K) and thalamus (L-N). (I, L) Quantifying average intersections of microglial processes across radial distances from the soma shows a significant reduction in the ABX group compared to VH in the cortex. (J, M) The average process length decreases in the ABX-treated group in the cortex (p = 0.04) but not in the thalamus. (K, N) Average number of branch nodes, with no significant differences observed between groups. (O-R) Representative images of in situ hybridization for pro-inflammatory cytokines IL-6 (O, P) and TNFα (Q, R) in microglia (Iba-1, green) with nuclear counterstain (DAPI, blue), showing increased expression in the VH-treated group compared to ABX. (S, T) IL-6 mRNA+ cell counts were significantly higher in the 2xTBI-VH group than in ABX in both the cortex and thalamus. (U, V) TNFα mRNA+ cell counts were also elevated in the cortex of the VH group (p < 0.01), though no significant differences were found in the thalamus. These results suggest that ABX treatment mitigates microglial activation and neuroinflammatory responses following repeated TBI, as evidenced by reduced microglial complexity and lower pro-inflammatory cytokine expression.

**Supplementary Figure 2.**
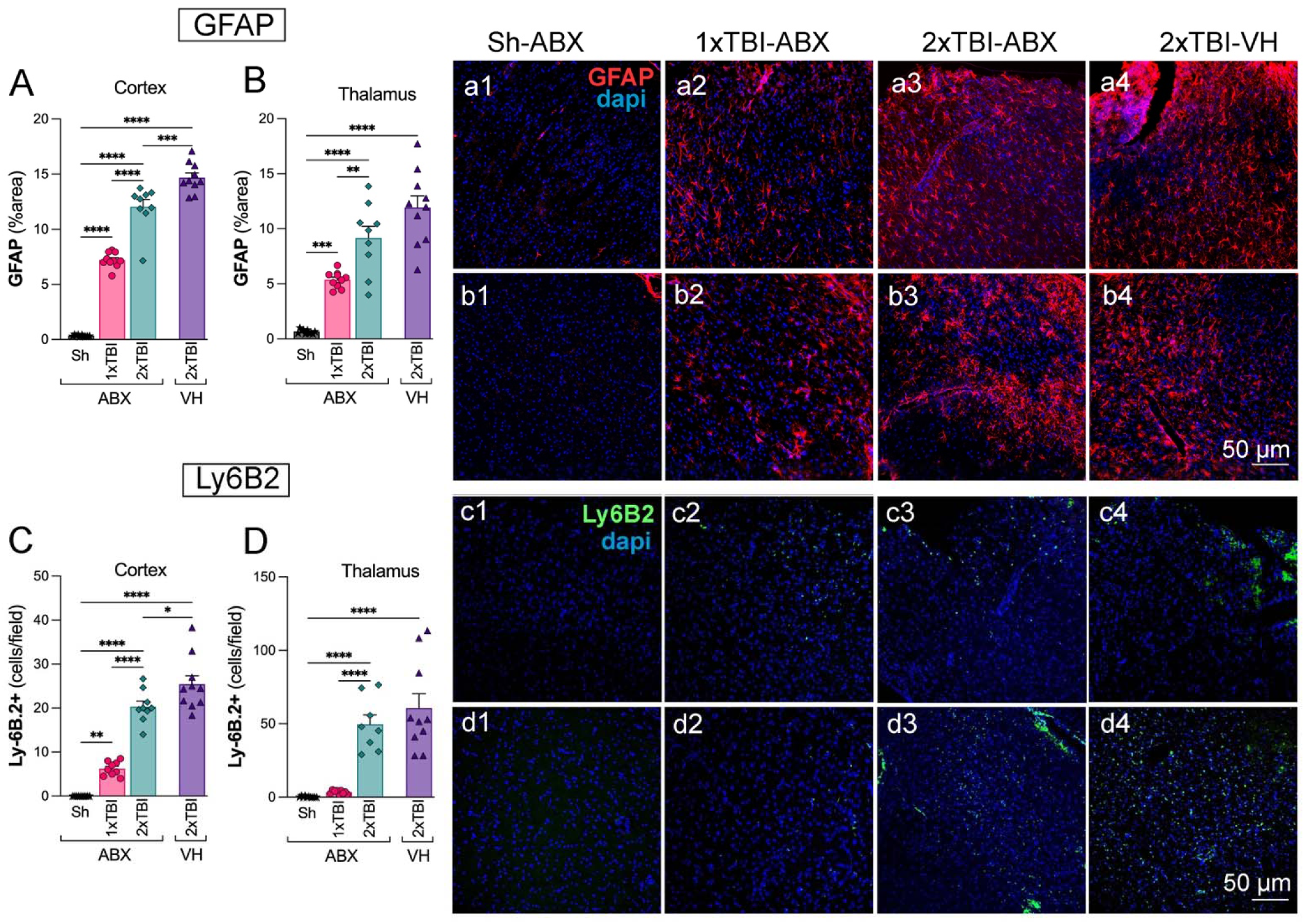
Antibiotic treatment decreases astrocyte reactivity but does not affect neutrophil infiltration following TBI. (A, B) Quantification of GFAP+ staining (% area) in the cortex (A) and thalamus (B) of Sh-ABX, 1xTBI-ABX, 2xTBI-ABX, and 2xTBI-VH groups. (a1–a4, b1– b4) Representative GFAP immunofluorescence images (red) with DAPI nuclear staining (blue) in the cortex (a1–a4) and thalamus (b1–b4) of the respective groups. (C, D) Quantification of Ly6B2+ immune cells (cells/field) in the cortex (C) and thalamus (D) of Sh-ABX, 1xTBI-ABX, 2xTBI-ABX, and 2× TBI-VH groups (c1–c4, d1–d4). Representative immunofluorescence images for Ly6B2 (green) with DAPI nuclear staining (blue) in the cortex (c1–c4) and thalamus (d1–d4) across groups. Statistical significance: *p < 0.05, **p < 0.01, ***p < 0.001, ****p < 0.0001 (one-way ANOVA with Tukey’s post hoc test) (n=10-9/group). Scale bar: 50 µm.

**Supplementary Figure 3.**
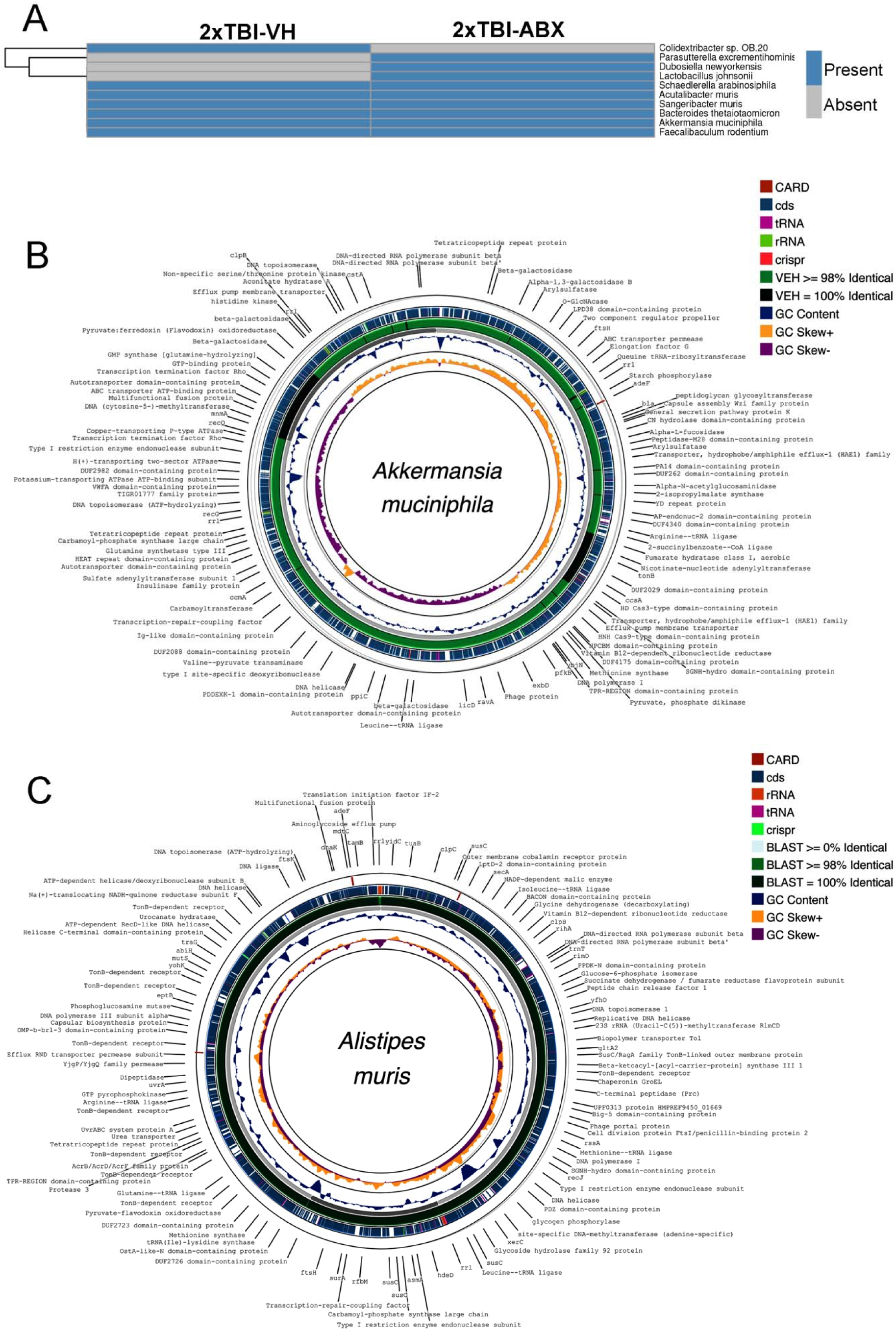
Metagenomic analysis reveals differential microbial composition and functional profiles in response to TBI and ABX treatment. (A) Presence-absence heatmap showing bacterial taxa identified in fecal samples from 2xTBI-VH and 2xTBI-ABX groups. Several bacterial species, including *Akkermansia muciniphila*, *Parasutterella excrementihominis*, and *Dubosiella newyorkensis*, were exclusively present or enriched in the VH-treated group. At the same time, other taxa, such as *Schaedlerella arabinosiphila* and *Sanguibacter muris*, were detected only in the ABX-treated group. (B) Circular genome visualization of *Akkermansia muciniphila*, highlighting annotated functional elements such as resistance genes (CARD), coding sequences (cds), transfer RNAs (tRNA), ribosomal RNAs (rRNA), and clustered regularly interspaced short palindromic repeats (CRISPR). The green rings indicate the genomic content comparison between the VH and ABX groups based on BLAST identity thresholds (≥0%, ≥98%, 100%). GC content and GC skew patterns are displayed in the innermost tracks, providing insights into potential functional adaptations under TBI conditions. (C) Circular genome representation of *Alistipes muris*, displaying key functional genes and genomic features as described for *Akkermansia muciniphila*, including transporters, antibiotic resistance genes, and metabolic regulators. Comparative genomic analysis reveals differences in gene presence and identity between the VH and ABX groups, suggesting functional adaptations in response to TBI and ABX treatment. These findings indicate distinct gut microbial compositions and genomic adaptations in response to TBI and antibiotic interventions, highlighting potential functional contributions of specific taxa to host-microbiome interactions post-injury.

## REFERENCES

1. Ng, S.Y., and Lee, A.Y.W. (2019). Traumatic Brain Injuries: Pathophysiology and Potential Therapeutic Targets. Front Cell Neurosci 13, 528. 10.3389/fncel.2019.00528.

2. Villapol, S., Byrnes, K.R., and Symes, A.J. (2014). Temporal dynamics of cerebral blood flow, cortical damage, apoptosis, astrocyte-vasculature interaction and astrogliosis in the pericontusional region after traumatic brain injury. Front Neurol 5, 82. 10.3389/fneur.2014.00082.

3. Veenith, T., Goon, S., and Burnstein, R.M. (2009). Molecular mechanisms of traumatic brain injury: the missing link in management. World J Emerg Surg 4, 7. 10.1186/1749-7922-4-7.

4. Jassam, Y.N., Izzy, S., Whalen, M., McGavern, D.B., and El Khoury, J. (2017). Neuroimmunology of Traumatic Brain Injury: Time for a Paradigm Shift. Neuron 95, 1246–1265. 10.1016/j.neuron.2017.07.010.

5. Kempuraj, D., Thangavel, R., Natteru, P.A., Selvakumar, G.P., Saeed, D., Zahoor, H., Zaheer, S., Iyer, S.S., and Zaheer, A. (2016). Neuroinflammation Induces Neurodegeneration. J Neurol Neurosurg Spine 1.

6. Bansal, V., Costantini, T., Kroll, L., Peterson, C., Loomis, W., Eliceiri, B., Baird, A., Wolf, P., and Coimbra, R. (2009). Traumatic brain injury and intestinal dysfunction: uncovering the neuro-enteric axis. J Neurotrauma 26, 1353–1359. 10.1089/neu.2008.0858.

7. Mou, Y., Du, Y., Zhou, L., Yue, J., Hu, X., Liu, Y., Chen, S., Lin, X., Zhang, G., Xiao, H., and Dong, B. (2022). Gut Microbiota Interact With the Brain Through Systemic Chronic Inflammation: Implications on Neuroinflammation, Neurodegeneration, and Aging. Front Immunol 13, 796288. 10.3389/fimmu.2022.796288.

8. Zhang, H., Chen, Y., Wang, Z., Xie, G., Liu, M., Yuan, B., Chai, H., Wang, W., and Cheng, P. (2022). Implications of Gut Microbiota in Neurodegenerative Diseases. Front Immunol 13, 785644. 10.3389/fimmu.2022.785644.

9. Chiu, L.S., and Anderton, R.S. (2023). The role of the microbiota-gut-brain axis in long-term neurodegenerative processes following traumatic brain injury. Eur J Neurosci 57, 400–418. 10.1111/ejn.15892.

10. Yuan, B., Lu, X.J., and Wu, Q. (2021). Gut Microbiota and Acute Central Nervous System Injury: A New Target for Therapeutic Intervention. Front Immunol 12, 800796. 10.3389/fimmu.2021.800796.

11. Hanscom, M., Loane, D.J., and Shea-Donohue, T. (2021). Brain-gut axis dysfunction in the pathogenesis of traumatic brain injury. J Clin Invest 131. 10.1172/JCI143777.

12. George, A.K., Behera, J., Homme, R.P., Tyagi, N., Tyagi, S.C., and Singh, M. (2021). Rebuilding Microbiome for Mitigating Traumatic Brain Injury: Importance of Restructuring the Gut-Microbiome-Brain Axis. Mol Neurobiol 58, 3614–3627. 10.1007/s12035-021-02357-2.

13. Iftikhar, P.M., Anwar, A., Saleem, S., Nasir, S., and Inayat, A. (2020). Traumatic brain injury causing intestinal dysfunction: A review. J Clin Neurosci 79, 237–240. 10.1016/j.jocn.2020.07.019.

14. Ma, E.L., Smith, A.D., Desai, N., Cheung, L., Hanscom, M., Stoica, B.A., Loane, D.J., Shea-Donohue, T., and Faden, A.I. (2017). Bidirectional brain-gut interactions and chronic pathological changes after traumatic brain injury in mice. Brain Behav Immun 66, 56–69. 10.1016/j.bbi.2017.06.018.

15. Erny, D., Hrabe de Angelis, A.L., Jaitin, D., Wieghofer, P., Staszewski, O., David, E., Keren-Shaul, H., Mahlakoiv, T., Jakobshagen, K., Buch, T., et al. (2015). Host microbiota constantly control maturation and function of microglia in the CNS. Nat Neurosci 18, 965–977. 10.1038/nn.4030.

16. Guo, C., Huo, Y.J., Li, Y., Han, Y., and Zhou, D. (2022). Gut-brain axis: Focus on gut metabolites short-chain fatty acids. World J Clin Cases 10, 1754–1763. 10.12998/wjcc.v10.i6.1754.

17. Overby, H.B., and Ferguson, J.F. (2021). Gut Microbiota-Derived Short-Chain Fatty Acids Facilitate Microbiota:Host Cross talk and Modulate Obesity and Hypertension. Curr Hypertens Rep 23, 8. 10.1007/s11906-020-01125-2.

18. Parada Venegas, D., De la Fuente, M.K., Landskron, G., Gonzalez, M.J., Quera, R., Dijkstra, G., Harmsen, H.J.M., Faber, K.N., and Hermoso, M.A. (2019). Short Chain Fatty Acids (SCFAs)-Mediated Gut Epithelial and Immune Regulation and Its Relevance for Inflammatory Bowel Diseases. Front Immunol 10, 277. 10.3389/fimmu.2019.00277.

19. Zhou, Y., Xu, H., Xu, J., Guo, X., Zhao, H., Chen, Y., Zhou, Y., and Nie, Y. (2021). F. prausnitzii and its supernatant increase SCFAs-producing bacteria to restore gut dysbiosis in TNBS-induced colitis. AMB Express 11, 33. 10.1186/s13568-021-01197-6.

20. Treangen, T.J., Wagner, J., Burns, M.P., and Villapol, S. (2018). Traumatic Brain Injury in Mice Induces Acute Bacterial Dysbiosis Within the Fecal Microbiome. Front Immunol 9, 2757. 10.3389/fimmu.2018.02757.

21. Howard, B.M., Kornblith, L.Z., Christie, S.A., Conroy, A.S., Nelson, M.F., Campion, E.M., Callcut, R.A., Calfee, C.S., Lamere, B.J., Fadrosh, D.W., et al. (2017). Characterizing the gut microbiome in trauma: significant changes in microbial diversity occur early after severe injury. Trauma Surg Acute Care Open 2, e000108. 10.1136/tsaco-2017-000108.

22. Ho, K.M., Kalgudi, S., Corbett, J.M., and Litton, E. (2020). Gut microbiota in surgical and critically ill patients. Anaesth Intensive Care 48, 179–195. 10.1177/0310057X20903732.

23. Dhillon, N.K., Adjamian, N., Fierro, N.M., Conde, G., Barmparas, G., and Ley, E.J. (2022). Early Antibiotic Administration is Independently Associated with Improved Survival in Traumatic Brain Injury. J Surg Res 270, 495–502. 10.1016/j.jss.2021.10.015.

24. Zhao, Q., Li, H., Li, H., and Zhang, J. (2023). Research progress on pleiotropic neuroprotective drugs for traumatic brain injury. Front Pharmacol 14, 1185533. 10.3389/fphar.2023.1185533.

25. Simon, D.W., Rogers, M.B., Gao, Y., Vincent, G., Firek, B.A., Janesko-Feldman, K., Vagni, V., Kochanek, P.M., Ozolek, J.A., Mollen, K.P., et al. (2020). Depletion of gut microbiota is associated with improved neurologic outcome following traumatic brain injury. Brain Res 1747, 147056. 10.1016/j.brainres.2020.147056.

26. Frohlich, E.E., Farzi, A., Mayerhofer, R., Reichmann, F., Jacan, A., Wagner, B., Zinser, E., Bordag, N., Magnes, C., Frohlich, E., et al. (2016). Cognitive impairment by antibiotic-induced gut dysbiosis: Analysis of gut microbiota-brain communication. Brain Behav Immun 56, 140–155. 10.1016/j.bbi.2016.02.020.

27. Dethlefsen, L., Huse, S., Sogin, M.L., and Relman, D.A. (2008). The pervasive effects of an antibiotic on the human gut microbiota, as revealed by deep 16S rRNA sequencing. PLoS Biol 6, e280. 10.1371/journal.pbio.0060280.

28. Faden, A.I., and Loane, D.J. (2015). Chronic neurodegeneration after traumatic brain injury: Alzheimer disease, chronic traumatic encephalopathy, or persistent neuroinflammation? Neurotherapeutics 12, 143–150. 10.1007/s13311-014-0319-5.

29. Aungst, S.L., Kabadi, S.V., Thompson, S.M., Stoica, B.A., and Faden, A.I. (2014). Repeated mild traumatic brain injury causes chronic neuroinflammation, changes in hippocampal synaptic plasticity, and associated cognitive deficits. J Cereb Blood Flow Metab 34, 1223–1232. 10.1038/jcbfm.2014.75.

30. Onyszchuk, G., LeVine, S.M., Brooks, W.M., and Berman, N.E. (2009). Post-acute pathological changes in the thalamus and internal capsule in aged mice following controlled cortical impact injury: a magnetic resonance imaging, iron histochemical, and glial immunohistochemical study. Neurosci Lett 452, 204–208. 10.1016/j.neulet.2009.01.049.

31. Zhao, S., Wang, X., Gao, X., and Chen, J. (2018). Delayed and progressive damages to juvenile mice after moderate traumatic brain injury. Sci Rep 8, 7339. 10.1038/s41598-018-25475-9.

32. Rogers, M.B., Simon, D., Firek, B., Silfies, L., Fabio, A., Bell, M.J., Yeh, A., Azar, J., Cheek, R., Kochanek, P.M., et al. (2022). Temporal and Spatial Changes in the Microbiome Following Pediatric Severe Traumatic Brain Injury. Pediatr Crit Care Med 23, 425–434. 10.1097/PCC.0000000000002929.

33. Bassetti, S., Tschudin-Sutter, S., Egli, A., and Osthoff, M. (2022). Optimizing antibiotic therapies to reduce the risk of bacterial resistance. Eur J Intern Med 99, 7–12. 10.1016/j.ejim.2022.01.029.

34. Dorner, P.J., Anandakumar, H., Rowekamp, I., Fiocca Vernengo, F., Millet Pascual-Leone, B., Krzanowski, M., Sellmaier, J., Bruning, U., Fritsche-Guenther, R., Pfannkuch, L., et al. (2024). Clinically used broad-spectrum antibiotics compromise inflammatory monocyte-dependent antibacterial defense in the lung. Nat Commun 15, 2788. 10.1038/s41467-024-47149-z.

35. Ramirez, J., Guarner, F., Bustos Fernandez, L., Maruy, A., Sdepanian, V.L., and Cohen, H. (2020). Antibiotics as Major Disruptors of Gut Microbiota. Front Cell Infect Microbiol 10, 572912. 10.3389/fcimb.2020.572912.

36. Pyles, R.B., Miller, A.L., Urban, R.J., Sheffield-Moore, M., Wright, T.J., Maxwell, C.A., Randolph, K.M., Danesi, C.P., McGovern, K.A., Vargas, J., et al. (2024). The altered TBI fecal microbiome is stable and functionally distinct. Front Mol Neurosci 17, 1341808. 10.3389/fnmol.2024.1341808.

37. Pelaseyed, T., Bergstrom, J.H., Gustafsson, J.K., Ermund, A., Birchenough, G.M., Schutte, A., van der Post, S., Svensson, F., Rodriguez-Pineiro, A.M., Nystrom, E.E., et al. (2014). The mucus and mucins of the goblet cells and enterocytes provide the first defense line of the gastrointestinal tract and interact with the immune system. Immunol Rev 260, 8–20. 10.1111/imr.12182.

38. Bercik, P., Denou, E., Collins, J., Jackson, W., Lu, J., Jury, J., Deng, Y., Blennerhassett, P., Macri, J., McCoy, K.D., et al. (2011). The intestinal microbiota affect central levels of brain-derived neurotropic factor and behavior in mice. Gastroenterology 141, 599–609, 609 e591-593. 10.1053/j.gastro.2011.04.052.

39. Nandwana, V., Nandwana, N.K., Das, Y., Saito, M., Panda, T., Das, S., Almaguel, F., Hosmane, N.S., and Das, B.C. (2022). The Role of Microbiome in Brain Development and Neurodegenerative Diseases. Molecules 27. 10.3390/molecules27113402.

40. Xie, X., Chen, X., Zhang, S., Liu, J., Zhang, W., and Cao, Y. (2024). Neutralizing gut-derived lipopolysaccharide as a novel therapeutic strategy for severe leptospirosis. Elife 13. 10.7554/eLife.96065.

41. Barrett, J.P., Costello, D.A., O’Sullivan, J., Cowley, T.R., and Lynch, M.A. (2015). Bone marrow-derived macrophages from aged rats are more responsive to inflammatory stimuli. J Neuroinflammation 12, 67. 10.1186/s12974-015-0287-7.

42. Glitza, I.C., Seo, Y.D., Spencer, C.N., Wortman, J.R., Burton, E.M., Alayli, F.A., Loo, C.P., Gautam, S., Damania, A., Densmore, J., et al. (2024). Randomized Placebo-Controlled, Biomarker-Stratified Phase Ib Microbiome Modulation in Melanoma: Impact of Antibiotic Preconditioning on Microbiome and Immunity. Cancer Discov 14, 1161–1175. 10.1158/2159-8290.CD-24-0066.

43. Gu, Y., Ye, T., Tan, P., Tong, L., Ji, J., Gu, Y., Shen, Z., Shen, X., Lu, X., and Huang, C. (2021). Tolerance-inducing effect and properties of innate immune stimulation on chronic stress-induced behavioral abnormalities in mice. Brain Behav Immun 91, 451–471. 10.1016/j.bbi.2020.11.002.

44. Drobyshevsky, A., Synowiec, S., Goussakov, I., Fabres, R., Lu, J., and Caplan, M. (2024). Intestinal microbiota modulates neuroinflammatory response and brain injury after neonatal hypoxia-ischemia. Gut Microbes 16, 2333808. 10.1080/19490976.2024.2333808.

45. Celorrio, M., Abellanas, M.A., Rhodes, J., Goodwin, V., Moritz, J., Vadivelu, S., Wang, L., Rodgers, R., Xiao, S., Anabayan, I., et al. (2021). Gut microbial dysbiosis after traumatic brain injury modulates the immune response and impairs neurogenesis. Acta Neuropathol Commun 9, 40. 10.1186/s40478-021-01137-2.

46. Lange, K., Buerger, M., Stallmach, A., and Bruns, T. (2016). Effects of Antibiotics on Gut Microbiota. Dig Dis 34, 260–268. 10.1159/000443360.

47. Zhang, S., and Chen, D.C. (2019). Facing a new challenge: the adverse effects of antibiotics on gut microbiota and host immunity. Chin Med J (Engl) 132, 1135–1138. 10.1097/CM9.0000000000000245.

48. Bicknell, B., Liebert, A., Borody, T., Herkes, G., McLachlan, C., and Kiat, H. (2023). Neurodegenerative and Neurodevelopmental Diseases and the Gut-Brain Axis: The Potential of Therapeutic Targeting of the Microbiome. Int J Mol Sci 24. 10.3390/ijms24119577.

49. Sgro, M., Iacono, G., Yamakawa, G.R., Kodila, Z.N., Marsland, B.J., and Mychasiuk, R. (2022). Age matters: Microbiome depletion prior to repeat mild traumatic brain injury differentially alters microbial composition and function in adolescent and adult rats. PLoS One 17, e0278259. 10.1371/journal.pone.0278259.

50. Peng, N., Wang, J., Zhu, H., Liu, Z., Ren, J., Li, W., and Wang, Y. (2024). Protective effect of carbon dots as antioxidants on intestinal inflammation by regulating oxidative stress and gut microbiota in nematodes and mouse models. Int Immunopharmacol 131, 111871. 10.1016/j.intimp.2024.111871.

51. Bravo, J.A., Forsythe, P., Chew, M.V., Escaravage, E., Savignac, H.M., Dinan, T.G., Bienenstock, J., and Cryan, J.F. (2011). Ingestion of Lactobacillus strain regulates emotional behavior and central GABA receptor expression in a mouse via the vagus nerve. Proc Natl Acad Sci U S A 108, 16050–16055. 10.1073/pnas.1102999108.

52. Bao, W., Sun, Y., Lin, Y., Yang, X., and Chen, Z. (2023). An integrated analysis of gut microbiota and the brain transcriptome reveals host-gut microbiota interactions following traumatic brain injury. Brain Res 1799, 148149. 10.1016/j.brainres.2022.148149.

53. Holcomb, M., Marshall, A., Flinn, H., Lozano, M., Soriano, S., Gomez-Pinilla, F., Treangen, T.J., and Villapol, S. (2024). Probiotic treatment causes sex-specific neuroprotection after traumatic brain injury in mice. Res Sq. 10.21203/rs.3.rs-4196801/v1.

54. Ma, Y., Liu, T., Fu, J., Fu, S., Hu, C., Sun, B., Fan, X., and Zhu, J. (2019). Lactobacillus acidophilus Exerts Neuroprotective Effects in Mice with Traumatic Brain Injury. J Nutr 149, 1543–1552. 10.1093/jn/nxz105.

55. Nagarajan, A., Scoggin, K., Gupta, J., Threadgill, D.W., and Andrews-Polymenis, H.L. (2023). Using the collaborative cross to identify the role of host genetics in defining the murine gut microbiome. Microbiome 11, 149. 10.1186/s40168-023-01552-8.

56. Otaru, N., Ye, K., Mujezinovic, D., Berchtold, L., Constancias, F., Cornejo, F.A., Krzystek, A., de Wouters, T., Braegger, C., Lacroix, C., and Pugin, B. (2021). GABA Production by Human Intestinal Bacteroides spp.: Prevalence, Regulation, and Role in Acid Stress Tolerance. Front Microbiol 12, 656895. 10.3389/fmicb.2021.656895.

57. Li, X., Zheng, S., Xu, H., Zhang, Z., Han, X., Wei, Y., Jin, H., Du, X., Xu, H., Li, M., et al. (2024). The direct and indirect inhibition of proinflammatory adipose tissue macrophages by acarbose in diet-induced obesity. Cell Rep Med, 101883. 10.1016/j.xcrm.2024.101883.

58. Soriano, S., Curry, K., Wang, Q., Chow, E., Treangen, T.J., and Villapol, S. (2022). Fecal Microbiota Transplantation Derived from Alzheimer’s Disease Mice Worsens Brain Trauma Outcomes in Wild-Type Controls. Int J Mol Sci 23. 10.3390/ijms23094476.

59. Guo, H., Pan, L., Li, L., Lu, J., Kwok, L., Menghe, B., Zhang, H., and Zhang, W. (2017). Characterization of Antibiotic Resistance Genes from Lactobacillus Isolated from Traditional Dairy Products. J Food Sci 82, 724–730. 10.1111/1750-3841.13645.

60. Dec, M., Nowaczek, A., Stepien-Pysniak, D., Wawrzykowski, J., and Urban-Chmiel, R. (2018). Identification and antibiotic susceptibility of lactobacilli isolated from turkeys. BMC Microbiol 18, 168. 10.1186/s12866-018-1269-6.

61. Jia, D.J., Wang, Q.W., Hu, Y.Y., He, J.M., Ge, Q.W., Qi, Y.D., Chen, L.Y., Zhang, Y., Fan, L.N., Lin, Y.F., et al. (2022). Lactobacillus johnsonii alleviates colitis by TLR1/2-STAT3 mediated CD206(+) macrophages(IL-10) activation. Gut Microbes 14, 2145843. 10.1080/19490976.2022.2145843.

62. Ding, Y.H., Qian, L.Y., Pang, J., Lin, J.Y., Xu, Q., Wang, L.H., Huang, D.S., and Zou, H. (2017). The regulation of immune cells by Lactobacilli: a potential therapeutic target for anti-atherosclerosis therapy. Oncotarget 8, 59915–59928. 10.18632/oncotarget.18346.

63. Xin, J., Zeng, D., Wang, H., Sun, N., Khalique, A., Zhao, Y., Wu, L., Pan, K., Jing, B., and Ni, X. (2020). Lactobacillus johnsonii BS15 improves intestinal environment against fluoride-induced memory impairment in mice-a study based on the gut-brain axis hypothesis. PeerJ 8, e10125. 10.7717/peerj.10125.

64. Tao, S., Fan, J., Li, J., Wu, Z., Yao, Y., Wang, Z., Wu, Y., Liu, X., Xiao, Y., and Wei, H. (2024). Extracellular vesicles derived from Lactobacillus johnsonii promote gut barrier homeostasis by enhancing M2 macrophage polarization. J Adv Res. 10.1016/j.jare.2024.03.011.

65. Straub, D., Blackwell, N., Langarica-Fuentes, A., Peltzer, A., Nahnsen, S., and Kleindienst, S. (2020). Interpretations of Environmental Microbial Community Studies Are Biased by the Selected 16S rRNA (Gene) Amplicon Sequencing Pipeline. Front Microbiol 11, 550420. 10.3389/fmicb.2020.550420.

66. McLaren, M.R., Willis, A.D., and Callahan, B.J. (2019). Consistent and correctable bias in metagenomic sequencing experiments. Elife 8. 10.7554/eLife.46923.

67. Villapol, S., Loane, D.J., and Burns, M.P. (2017). Sexual dimorphism in the inflammatory response to traumatic brain injury. Glia 65, 1423–1438. 10.1002/glia.23171.

68. Keane, S.P., Chadman, K.K., Gomez, A.R., and Hu, W. (2024). Pros and cons of narrow-versus wide-compartment rotarod apparatus: An experimental study in mice. Behav Brain Res 463, 114901. 10.1016/j.bbr.2024.114901.

69. Shan, H.M., Maurer, M.A., and Schwab, M.E. (2023). Four-parameter analysis in modified Rotarod test for detecting minor motor deficits in mice. BMC Biol 21, 177. 10.1186/s12915-023-01679-y.

70. Villapol, S., Balarezo, M.G., Affram, K., Saavedra, J.M., and Symes, A.J. (2015). Neurorestoration after traumatic brain injury through angiotensin II receptor blockage. Brain 138, 3299–3315. 10.1093/brain/awv172.

71. Caporaso, J.G., Lauber, C.L., Walters, W.A., Berg-Lyons, D., Huntley, J., Fierer, N., Owens, S.M., Betley, J., Fraser, L., Bauer, M., et al. (2012). Ultra-high-throughput microbial community analysis on the Illumina HiSeq and MiSeq platforms. ISME J 6, 1621–1624. 10.1038/ismej.2012.8.

72. Weisburg, W.G., Barns, S.M., Pelletier, D.A., and Lane, D.J. (1991). 16S ribosomal DNA amplification for phylogenetic study. J Bacteriol 173, 697–703. 10.1128/jb.173.2.697-703.1991.

73. McDonald, D., Jiang, Y., Balaban, M., Cantrell, K., Zhu, Q., Gonzalez, A., Morton, J.T., Nicolaou, G., Parks, D.H., Karst, S.M., et al. (2024). Greengenes2 unifies microbial data in a single reference tree. Nat Biotechnol 42, 715–718. 10.1038/s41587-023-01845-1.

74. Constantinides, B., Hunt, M., and Crook, D.W. (2023). Hostile: accurate decontamination of microbial host sequences. Bioinformatics 39. 10.1093/bioinformatics/btad728.

75. De Coster, W., and Rademakers, R. (2023). NanoPack2: population-scale evaluation of long-read sequencing data. Bioinformatics 39. 10.1093/bioinformatics/btad311.

76. Li, H., Handsaker, B., Wysoker, A., Fennell, T., Ruan, J., Homer, N., Marth, G., Abecasis, G., Durbin, R., and Genome Project Data Processing, S. (2009). The Sequence Alignment/Map format and SAMtools. Bioinformatics 25, 2078–2079. 10.1093/bioinformatics/btp352.

77. Chaumeil, P.A., Mussig, A.J., Hugenholtz, P., and Parks, D.H. (2019). GTDB-Tk: a toolkit to classify genomes with the Genome Taxonomy Database. Bioinformatics 36, 1925–1927. 10.1093/bioinformatics/btz848.

78. Chklovski, A., Parks, D.H., Woodcroft, B.J., and Tyson, G.W. (2023). CheckM2: a rapid, scalable and accurate tool for assessing microbial genome quality using machine learning. Nat Methods 20, 1203–1212. 10.1038/s41592-023-01940-w.

79. Li, H. (2018). Minimap2: pairwise alignment for nucleotide sequences. Bioinformatics 34, 3094–3100. 10.1093/bioinformatics/bty191.

80. Nissen, J.N., Johansen, J., Allesoe, R.L., Sonderby, C.K., Armenteros, J.J.A., Gronbech, C.H., Jensen, L.J., Nielsen, H.B., Petersen, T.N., Winther, O., and Rasmussen, S. (2021). Improved metagenome binning and assembly using deep variational autoencoders. Nat Biotechnol 39, 555–560. 10.1038/s41587-020-00777-4.

81. Pan, S., Zhao, X.M., and Coelho, L.P. (2023). SemiBin2: self-supervised contrastive learning leads to better MAGs for short- and long-read sequencing. Bioinformatics 39, i21–i29. 10.1093/bioinformatics/btad209.

82. Sieber, C.M.K., Probst, A.J., Sharrar, A., Thomas, B.C., Hess, M., Tringe, S.G., and Banfield, J.F. (2018). Recovery of genomes from metagenomes via a dereplication, aggregation and scoring strategy. Nat Microbiol 3, 836–843. 10.1038/s41564-018-0171-1.

83. Wu, Y.W., Simmons, B.A., and Singer, S.W. (2016). MaxBin 2.0: an automated binning algorithm to recover genomes from multiple metagenomic datasets. Bioinformatics 32, 605–607. 10.1093/bioinformatics/btv638.

84. Balaji, A., Kille, B., Kappell, A.D., Godbold, G.D., Diep, M., Elworth, R.A.L., Qian, Z., Albin, D., Nasko, D.J., Shah, N., et al. (2022). SeqScreen: accurate and sensitive functional screening of pathogenic sequences via ensemble learning. Genome Biol 23, 133. 10.1186/s13059-022-02695-x.

85. Sapoval, N., Liu, Y., Curry, K.D., Kille, B., Huang, W., Kokroko, N., Nute, M.G., Tyshaieva, A., Dilthey, A., Molloy, E.K., and Treangen, T.J. (2024). Lightweight taxonomic profiling of long-read metagenomic datasets with Lemur and Magnet. bioRxiv. 10.1101/2024.06.01.596961.

86. Zinger, A., Soriano, S., Baudo, G., De Rosa, E., Taraballi, F., and Villapol, S. (2021). Biomimetic Nanoparticles as a Theranostic Tool for Traumatic Brain Injury. Adv Funct Mater 31, 2100722. 10.1002/adfm.202100722.

87. Fe Lanfranco, M., Loane, D.J., Mocchetti, I., Burns, M.P., and Villapol, S. (2017). Combination of Fluorescent in situ Hybridization (FISH) and Immunofluorescence Imaging for Detection of Cytokine Expression in Microglia/Macrophage Cells. Bio Protoc 7. 10.21769/BioProtoc.2608.

